# Neural Representations of Extrasystoles: A Predictive Coding Perspective

**DOI:** 10.1101/2024.09.06.610728

**Authors:** Pia Reinfeld, Tim Paul Steinfath, Pei-Hsin Ku, Vadim Nikulin, Jane Neumann, Arno Villringer

## Abstract

The coordinated interplay between brain and body is essential for survival, with the heart-brain axis playing a fundamental role in perceptual, cognitive and motor processes. Central to this interaction, the heart-evoked potential (HEP) represents a neural correlate of the heart activity. Further, the HEP is thought to represent a neuronal prediction for each heartbeat, raising questions about its role in arrhythmia. Yet, previous studies have primarily focused on regular heart rhythms, with only a few investigations delving into arrhythmias and, notably, none into extrasystoles, a form of benign arrhythmias that disrupt the regular heart rhythm. Extrasystoles represent a unique possibility for investigating brain responses to unexpected pronounced alterations in heart rhythm. We took advantage of the largest dataset (103 subjects with 3000 extrasystoles), including both EEG and ECG recordings, and analysed the neural response to both types of extrasystoles (supraventricular and ventricular) using multiverse approaches and control analysis in time and source space. We found that the HEP was significantly reduced for ventricular extrasystoles with underlying sources in the left insular. For the postextrasystolic beat of both types of extrasystoles, we found a significantly earlier and increased HEP originating from sources in the left frontal orbital cortex and the anterior cingulate gyri. The reduced HEP response to the ventricular extrasystole might result from inactive interoceptive cardiac pathways. In contrast, increased HEP of the postextrasystolic beat along with the anatomical neural HEP generators may reflect an interoceptive prediction error arising from a naturally occurring discrepancy between the predicted and actual heart rhythm, with a main source in the anterior cingulate gyri, a primary location for prediction error processing.

## 1 Introduction

The interaction between brain and body is essential for survival. A key axis is the heart-brain interaction. Within this axis, emotions like fear, primarily originating from the brain, increase heart activity, preparing the body for a potential escape. Conversely, the heart also influences the brain, affecting cognition and emotion.

The HEP reflects the cortical processing of cardiac signals. It can be obtained by averaging electroencephalography (EEG) signals that are time-locked to the R- or T-peaks of heartbeats measured by electrocardiography (ECG). However, the exact mechanisms behind the generation of HEPs still need to be fully understood. [1]

HEPs have been contextualised with emotion, self and consciousness and within the predictive coding framework. [2] Within the latter, the brain constantly generates and updates a model of the environment to predict what might happen next and when. A prediction error occurs if the prediction does not match the sensory input, which might be reflected in a mismatch negativity (MMN) response. [3] Prediction errors manifest in HEPs in tasks where the heartbeat is coupled to exteroceptive stimuli. Pfeiffer et al. found a surprise response in HEPs when an expected sound was omitted from a sequence of sounds synchronised to the heartbeat. [4, 5] This predictive mechanism extends beyond exteroceptive anticipation to include the interoceptive sensory stimulus of the regular heartbeat. It is thought that the brain interprets the interoceptive stimulus of the regular heartbeat as a predictable event [6]. However, predictive coding modalities have traditionally been tested in somewhat artificial setups, where experiments are designed with controlled changes to stimuli or signals. To date, no study has investigated predictive coding using naturally occuring stimuli, specifically in the context of interoceptive prediction errors, stimuli within the heart. A prime example of such a natural prediction error could be a premature heart-beat occuring outside the physiological heart rhythm, known as extrasystoles (ES).

ES are arrhythmias affecting affecting about 60 % of the healthy population [7]. They occur predominantly idiopathically and are subclassified into supraventricular (SVES) and ventricular extrasystoles (VES).

VESs are induced by early ventricular depolarisation, leading to premature and abnormal QRS-complexes. They manifest polymorphically, indicating varied origins of the electrical impulses within the ventricles. VESs typically cause reduced stroke volume due to the short inter-beat interval (IBI), asymmetric contraction of the ventricles, and an absence of presystolic atrial contraction [8]. VES frequency scales with growing age, higher blood pressure, smoking, or less physical activity, while the fundamental cause of VESs is still largely unknown [9].

SVESs originate in the atria myocardium, leading to a contraction of the atria and mostly appear with normal premature QRS-complex. They lead to less abnormal hemodynamics than VESs due to a remaining presystolic atrial contraction. [8] Higher incidences of SVESs are associated with many causes, including medical pathologies like hypertension and diabetes mellitus or structural causes like coronary artery disease [10].

A high frequency of ESs is associated with an increased risk of new-onset atrial fibrillation, ischemic strokes, and cardiomyopathy [9, 11, 10].

They appear commonly asymptomatic. However, symptoms like palpitations, fatigue, shortness of breath and, anxiety may appear [10, 9].

Since ESs are premature heartbeats that occur outside the regular heart rhythm, we hypothesised that these irregular beats can be perceived as disruptions in the expected pattern of heartbeats. Furthermore, according to the predictive coding theory [5, 12], the brain continuously generates predictions about the timing and regularity of heartbeats based on past experiences. Therefore, when an ES occurs, it could deviate from the predicted pattern, resulting in a significant prediction error, which, in turn, can be reflected in a larger HEP amplitude. Here, we are specifically interested in the time window of the extrasystole and the postextrasystolic beat, as those are the beats often reported to be perceived [9].

Other studies looking into different types of arrhythmias found modulatory effects on the neural representations. For example, Kumral et al. have shown that the HEP is reduced in patients with atrial fibrillation [13].

Alternatively, the HEP during ESs could reflect the same impaired heart-brain interaction as seen in atrial fibrillation, given that both conditions are types of arrhythmias. This attenuated interoception could also manifest as a diminished HEP amplitude. [13].

To test these hypotheses, we compared HEPs belonging to more than 3000 ESs of 103 participants to their corresponding normal heartbeats, with a particular emphasis on the beat after the ES, as this is the beat that is often reported to be perceived [9]. We used a multiverse pipeline to analyse the EEG data to ensure the robustness of the results against parameter variations. Traditionally, research findings are presented based on a limited number of analysis pipelines. The multiverse analysis method expands this by evaluating a variety of reasonable pipelines and reporting outcomes across all of them. This strategy examines the variability in results and evaluates how the selection of different pipelines affects the robustness and reliability of the observed findings. We further investigated the potential differences between neuronal responses to SVESs and VESs. To reveal the underlying sources, we conducted a multiverse source analysis. Correlation analyses between the HEP and ECG amplitude differences and a control analysis for IBI and T-peak amplitude were performed to assess the effect of possible residual heart-beat and cardiac field artefacts (CFA), which is an electric field generated by the heart spreading into the scalp electrodes.

## 2 Methodologies

### 2.1 Participants

Participants were selected from the population-based study of the Leipzig Research Centre for Civilization Diseases (LIFE-Adult). [14] Initially, we selected 3492 participants who had simultaneously recorded ECG and resting state EEG. Out of 2228 participants who had at least one SVES or VES beat, 103 participants meeting the criteria were enrolled for analysis, including 51 for the SVES condition and 52 for the VES condition (Supplementary Figure 10). We only included participants with at least 25 isolated SVES or VES beats. To minimise the influence of potential cardiac problems, participants with more than five detected SVES/VES trigeminy, bigeminy, run, couplet, prolonged R-R interval, tachycardia, fibrillation, or flutter were excluded from the study. All the ECG patterns were first automatically detected by the CER-S (Continuous ECG Recording Suite) software developed by AMPS (Analysing Medical Parameters for Solutions). CER-S is designed for scientific ECG analysis and supports data formats from 1- to 12-channel ECGs and open-standard formats like MIT-WFDB, which was utilised in this study. [15]. The software employs the ABILE algorithm to detect and analyse heartbeats in an ECG. For this study, the maximum coupling interval for detecting ESs was set to 75 %. After detection, two researchers manually inspected and finalised the beats as described in Section 2.3. No participants included in this study were using pacemakers. The mean age was 70.91 ± 4.51 years for people allocated in the SVES condition (male: 32, female: 19) and 70.76 ± 4.89 years for those in the VES condition (male: 33, female: 19).

### 2.2 Recordings

The 20 min resting EEG was recorded in an electrically shielded and sound-attenuated booth. 34 passive Ag/AgCl electrodes were used, including 31 scalp electrodes placed according to the 10 −20 system on an EEG cap. Additionally, two electrodes captured vertical and horizontal eye movements, while a bipolar electrode on the forearm simultaneously recorded the ECG. The signals were amplified using a QuickAmp amplifier (Brain Products). EEG signals were referenced against the common average and recorded at a sampling rate of 1000 Hz, with a low-pass filter set at 280 Hz. Impedance levels were kept below 10 kΩ.

### 2.3 ECG Preprocessing

We analysed all ECGs from participants with ECG and EEG recordings using CER-S along with custom-written bash scripts. The number of ESs for both types (SVES and VES) was extracted to select participants. We visually inspected the detected ESs and normal sinus beats within each selected participant. If necessary, we adjusted the detection trigger, which marked the timepoint of the R-peak, or excluded participants from the analysis if (1) the researchers were not able to identify a clear P-wave, (2) atrial fibrillation or flutter was present, or (3) the artefacts from the recordings have interfered the identification of cardiac rhythm. Due to the distinct morphology of VESs compared to normal heartbeats, the highest initial deflection was used as the ‘R-peak’ for VES detection.

We then imported the triggers for R-peaks of ESs and all other beats as events into EEGLAB/FieldTrip. ESs and sinus beats occurring in segments with EEG artefacts were excluded from the analysis. Isolated ESs, those with only sinus beats in the four beats preceding and five following the ES, were labelled accordingly. When the SVES and VES groups were combined, the events were labelled as ‘ES’ for the extrasystole itself. The additional labels ‘ES-1’, ‘ES-2’, and ‘ES-3’ were used for the beats preceding the extrasystole, and ‘ES+1’ and ‘ES+2’ for the beats following the extrasystole. This labelling scheme was similarly applied to both SVES and VES (Supplementary Figure 1). Regular sinus beats without any temporal relation to the extrasystoles were labelled as ‘N’.

In the recruited 103 participants, we detected 3202 ESs out of 51 participants for the SVES condition and 3345 ESs out of 52 for the VES condition. On average, there were 62.78 ± 50.24 ESs per person for SVES (range [26, 234]) and 64.33 ± 47.06 for VES (range [25, 238]). After the ECG preprocessing, 1443 SVESs and 1622 VESs met the criteria and were selected for analysis. This resulted in an average number of ESs per person of 28.86 ± 16.64 for SVES (range [8, 105]) and 31.19 ± 15.03 for VES (range [9, 77]).

### 2.4 EEG Preprocessing

The EEG was preprocessed using custom scripts and the MATLAB toolboxes EEGLAB and FieldTrip [16, 17, 6, 18]. For time domain analysis, the EEG data were filtered between 0.5 Hz and 20 Hz using a Hamming windowed sinc FIR filter. The filter order for the FIR filter was automatically determined as implemented in the pop_eegfiltnew function from EEGLAB [16]. The data was downsampled to 500 Hz, and events within segments exceeding 80 mV were excluded after visual inspection. The ECG events found by CER-S were added to the ECG and EEG signals as event markers. Extended Infomax algorithm-based independent component analysis removed artefacts from ocular, muscular, and heartbeat sources. To identify CFAs, the independent component (IC) and ECG time courses were segmented into intervals between −200 ms and −200 ms around the R-peaks and averaged across epochs. Next, average IC and ECG time courses were correlated, and ICs with a correlation greater than 0.8 with the ECG were marked as artefacts and manually inspected before rejection [19]. On average, 1.36 ± 0.8615 (range [0, 4]) cardiac field components were removed. Further settings were used to test the results’ robustness against variation in the preprocessing pipeline (Figure 4).

### 2.5 HEP

The EEG signals were segmented into epochs between −200 ms and 800 ms around the R-peaks and averaged to analyse the HEP in the time domain. To alleviate the impact of the ES-1 beat on ES due to the shortened IBI (compare Supplementary Tables 1 and 2), the averaged N beat was subtracted from each ES-1 beat before averaging. Baseline correction for all beats was performed using the time window from −150 ms to −50 ms prior to the reference condition ES-3 as baseline. This ensured that slow fluctuations did not impact the analysed HEP.

### 2.6 Reference Condition

The third beat before the extrasystole (ES-3) was used as the reference condition. This choice ensured an equivalent number of beats between the reference condition and the condition of interest (e.g., ES, ES+1), thereby maintaining a consistent signal-to-noise ratio (SNR) and increasing statistical robustness. This further ensures that, spontaneous brain activity occurring around the time of the ES is equally represented in both conditions, thereby preventing it from influencing the result.

### 2.7 Statistical Analysis

Cluster-based permutation *t*-tests were used to assess within-participant statistical significance between the conditions (ES-3 and ES/ES+1) using the FieldTrip toolbox. [20, 21] This method involves the clustering of samples whose *t*-values exceed a threshold (cluster threshold *p*-value: 0.05 two-sided) based on temporal, spatial (and spectral) adjacency and then calculating the cluster statistics by summing the *t*-values within each cluster. The Monte Carlo method was used to calculate the probability of significance while dealing with the problem of multiple comparisons by randomly permuting the condition labels 5000 times. Clusters with *p*-values less than 0.05 (corrected for a two-tailed test) and at least two adjacent electrodes in the neighbouring time frame were considered significant. The statistical analysis was performed on all EEG electrodes and the entire time window (−200 ms to 800 ms). A similar cluster-based permutation test was applied to the corresponding ECG signals.

### 2.8 Source Analysis

The source analysis was performed participant-wise using the ‘New York Head’ forward model [22]. This finite element method is based on the ICBM152 anatomical template [23]. The lead field matrix of 5004 source dipoles with fixed orientations perpendicular to the cortical surface was used. After removing channels PO5 and PO6, as they were not included in the pre-computed lead field, the lead field was transformed to the common mean reference. To solve the inverse problem, exact low-resolution tomography (eLORETA) [24] was applied using the METH toolbox (Guido Nolte; RRID: SCR_016104) with minor modifications by [25]. The regularisation parameter was varied between 0.001, 0.05, and 0.5 to assess the robustness of the results to parameter variations.

Furthermore, standard averaging and averaging with sign-flip were compared to aggregate the dipole sources into single regions of interest, as done by [26]. This multiverse source analysis was applied at the averaged time window where the main difference between conditions was found. To investigate the differences between conditions at the source level, ten regions of interest (ROIs) typically associated with HEP modulations were selected for both hemispheres: posterior cingulate gyrus, anterior cingulate gyrus (ACC), postcentral gyrus, frontal orbital cortex, and insular cortex; similar to [13]. A dependent *t*-test was applied to the participant-wise absolute data and corrected for False Discovery Rate (FDR).

### 2.9 Controlling for CFA: Correlation Analysis

To assess the effect of possible residual heartbeats and CFAs on the differences between the two conditions, we performed similar correlation analyses as Buot et al. [27] between the ECG and EEG.

#### Cluster-Based Permutation Analysis of entire Time Window

For all time courses and each participant, the epoch-wise difference (Δ) between the conditions was calculated. The empirical Spearman correlation between each averaged Δ EEG channel and the averaged Δ ECG channel for each time point of the entire time window and each participant was then determined and Fisher transformed. To estimate the correlation at the chance level, the associations between time points were permuted (e.g., time point *i* in the time course of electrode Fp1 was reallocated to time point *j* in the ECG-time course) and the correlation coefficient recalculated. After performing this permutation and recalculation 100 times, the median of the correlation coefficients was taken and statistically compared with the empirical correlation using a cluster-based permutation test at the group level.

#### Analysis of Averaged Time Window

In addition, a second correlation analysis focused on the time window in which the primary difference between conditions was identified in the EEG. This time window of the EEG and ECG signals was selected and averaged over time and then over epochs. The empirical Pearson correlation coefficients across participants and electrodes between the ECG and all EEG channels were calculated and corrected for FDR.

### 2.10 Control Analyses

We conducted two control analyses to ensure that the observed differences in ES+1 in the EEG were not due to variations in the ECG. These analyses involved matching epochs based on IBI and T-peak amplitude.

To determine the IBI of ES+1, we calculated the time between the R-peak of ES+1 and ES+2. We then searched for matching IBIs within N and ES-3 to find sufficient partners. The IBIs for these beats were computed as the time between the R-peak of each beat and the preceding beat.

To match the T-peak amplitude, we extracted the maximum absolute value for ES+1 within the 200 ms to 400 ms time window and matched it with the corresponding N and ES-3 beat values.

We solved the linear assignment problem with a predefined cost for unmatched epochs. Trials within ES+1 that did not have a counterpart were left unmatched and excluded from further analysis. The cost was iteratively adapted and adjusted until there was no significant difference in the dependent *t*-test between ES+1 values (T-peak amplitude/IBI) and the matched N/ES-3 values. After that, the cluster-based permutation *t*-tests were repeated.

## 3 Results

### 3.1 HEP Analysis

We analysed the impact of ESs on brain activity by examining the time windows of the ES and surrounding beats (ES-3, ES-2, ES-1, ES, ES+1, ES+2).

First, we confirmed that the third beat preceding the ES served as a suitable reference condition. The cluster-based permutation test was performed between ES-3 and N over the entire time window (−200 ms to 800 ms). This analysis showed no significant difference between the two beats (*p* > 0.05, Supplementary Figure 11), confirming the suitability of ES-3 as the reference condition.

The other HEPs of the preceding beats (ES-2 and ES-1) did not show a significant difference to the reference condition ES-3 (*p* > 0.05, Supplementary Figures 12 and 13). Further, no significant difference was observed between ES+2 (SVES+2, VES+2) and the reference condition (*p* > 0.05), suggesting a return to regular brain activity two beats after the ES (Supplementary Figure 14). Consequently, only the conditions with significant differences will be further discussed.

#### N

We analysed the grand average HEP for a normal heartbeat (N) over all 103 participants. The HEP begins around 200 ms after the R-peak and continues until 600 ms, which is consistent with the literature [28]. It appears bipolar, with a negative amplitude in the frontal-central electrodes and a positive in the occipital-central electrodes (Supplementary Figure 15).

#### ES

A cluster-based permutation test identified a significant difference between the ES and reference conditions. This difference was characterised by a diminished HEP in the ES condition, reflected in both a negative and a positive cluster around the 230 ms to 330 ms time window. The negative cluster was identified over central occipital electrodes (C3, C4, CP5, CP6, TP9, P3, P4, P7, P8, O1, O2, PO9, PO10; Monte Carlo *p* = 0.0132). The positive cluster was located over central-frontal electrodes (Fp1, Fp2, F3, F4, F7, F8, Fz, FC1, FC2, FC5, T7, FT9; Monte Carlo *p* = 0.0002).

The VES and SVES conditions were retested individually to determine whether the results were mainly driven by one type of ES.

#### VES

Performing the cluster-based permutation tests on the entire time window revealed a significant difference between the VES condition and the reference condition (VES-3) (*p* < 0.05) similar to the effect found in the ES condition. This corresponded to one negative and one positive cluster in the data from 220 ms to 350 ms. The negative cluster is located over central-occipital electrodes (C3, CP5, CP6, TP9, TP10, P3, P4, P7, P8, O1, O2, PO9, PO1; Monte Carlo *p* < 0.0002), whereas the positive cluster is found over central-frontal electrodes (Fp1, Fp2, F3, F4, F7, F8, Fz, FC1, FC5, FC6, C3, T7, FT9; Monte Carlo *p* = 0.0008, Figure 2).

**Figure 1:**
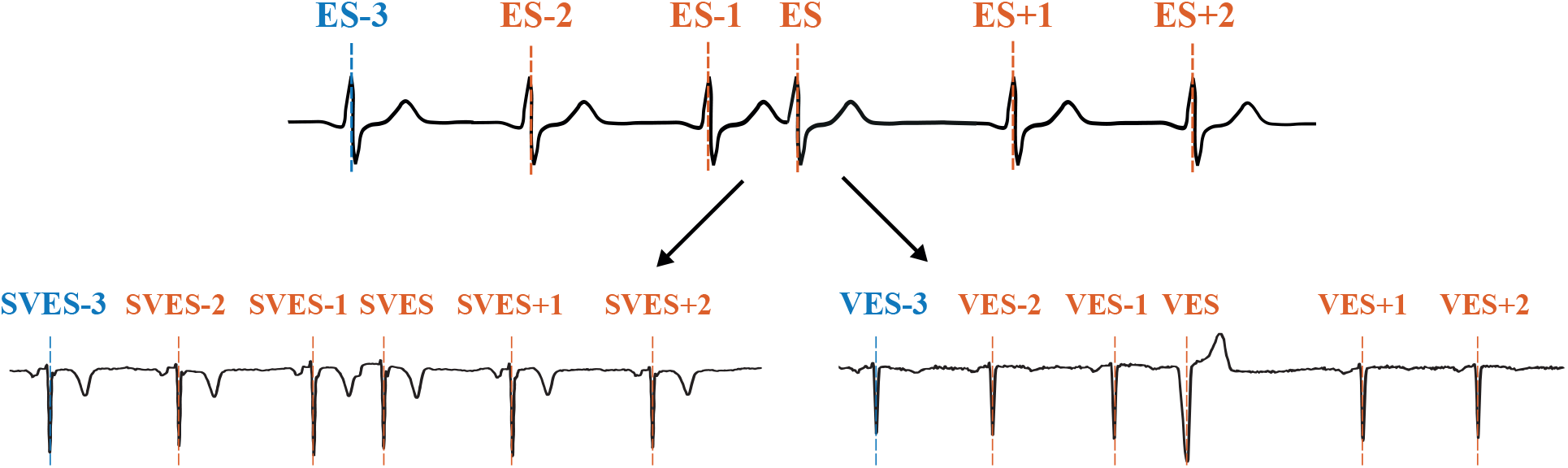
Illustration of the different beat types. The top row displays a schematic ECG where both types of ESs are aggregated. Each R-peak is annotated with the beat annotations used in this study. The bottom row shows ECGs derived from actual participant data for SVESs and VESs separately with beat annotations. Blue annotated the reference condition (compare Section 2.6).

**Figure 2:**
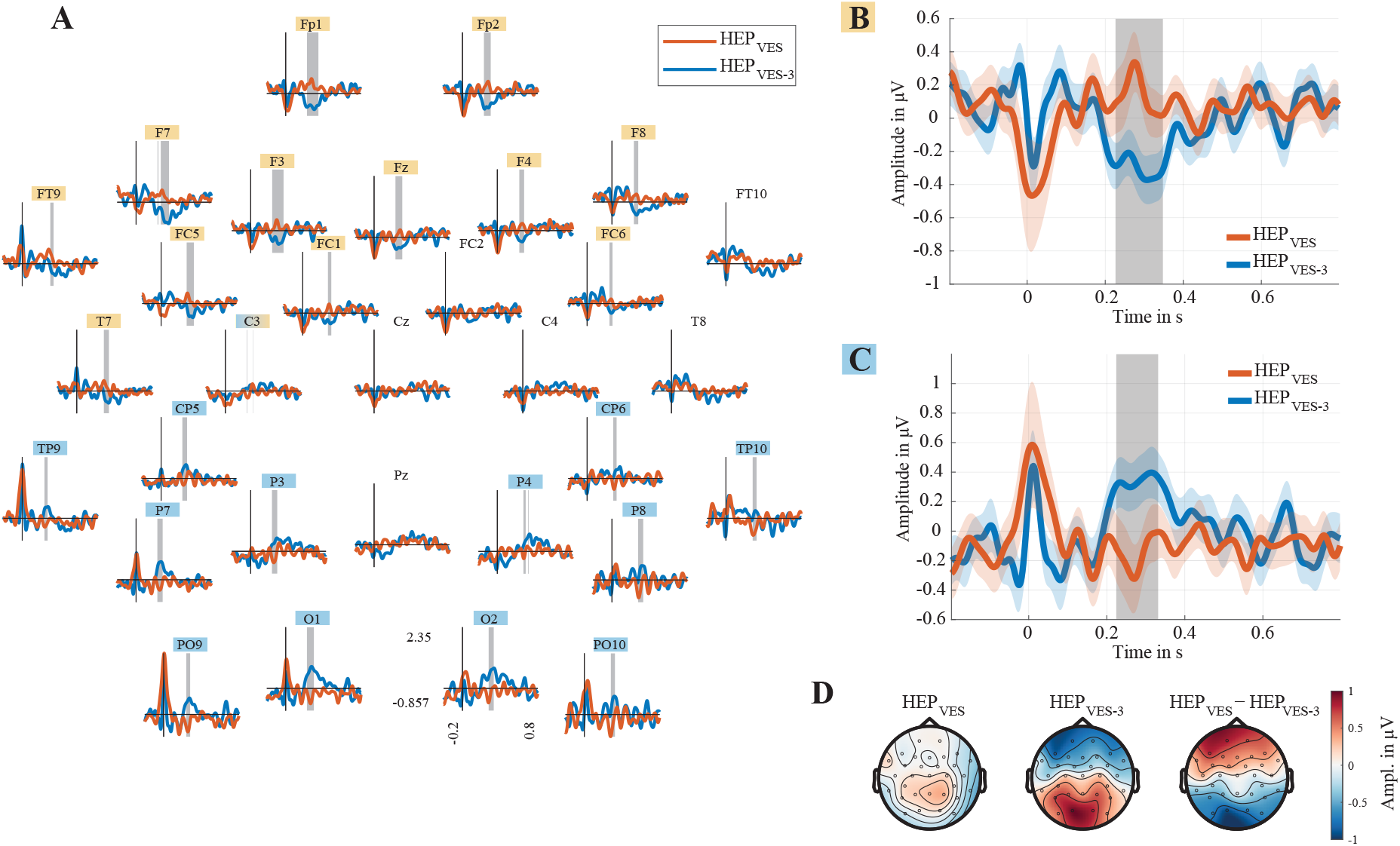
HEP of VES and reference condition (VES-3). Grand average over all VES participants. **(A)** HEPs in each EEG channel. **(B)** Positive Cluster averaged over central-frontal electrodes (Monte Carlo p = 0.0008) between the VES and reference conditions. Electrodes used for averaging are highlighted in orange. The grey-shaded area highlights the time window of the found cluster. The coloured-shaded regions around the lines mark the ±1 SEM. **(C)** Negative Cluster averaged over central-frontal electrodes (Monte Carlo p = 0.0002) between VES and reference condition. Electrodes used for averaging are highlighted in blue. The grey-shaded area highlights the time window of the found cluster. The coloured-shaded regions around the lines mark the ±1 SEM. **(D)** Topographies of HEPs in VES, VES-3 and differences between conditions between 220 ms to 350 ms after R-peaks.

Given the polymorphic nature of the VES (Supplementary Figure 16), averaging positive and negative polarities of the ‘T-wave’ might result in a flattend HEP due to the CFA. To ascertain, that averaging the ECG did not impact on the corresponding HEP, the analysis was reconducted separately on subsets of participants categorised by the polarity of the ‘T-wave’ of their ECG signals. Analyses of both subsets confirmed the initial finding and showed a significant difference between conditions (*p* < 0.05) with similar clusters.

#### SVES

No significant effect was observed for the analysis of SVES only (*p* > 0.05, Supplementary Figure 17). Thus, the effects observed for all ES was primarily driven by VES. Consecquently, further analyses, e.g. in source space, were restricted to the VES condition.

#### ES+1

Cluster-based permutation tests were further used to test whether the HEP of the postextrasystolic beat (ES+1) condition also differed from the HEP of the reference condition (ES-3). The analysis revealed a significant difference between the conditions (*p* < 0.05) corresponding to one positive and one negative cluster in the observed data from 130 ms to 200 ms (both Monte Carlo *p* = 0.002). The negative cluster is distributed over central-frontal (Fp1, Fp2, F3, F4, F7, F8, Fz, FC1, FC2, FC6, C4, Cz) and the positive cluster over central-occipital electrodes (CP5, CP6, TP9, TP10, P3, P4, P7, P8, O1, O2, PO9, PO10). Overall, the HEP appeared earlier and with increased amplitude in the ES+1 condition (Figure 3).

**Figure 3:**
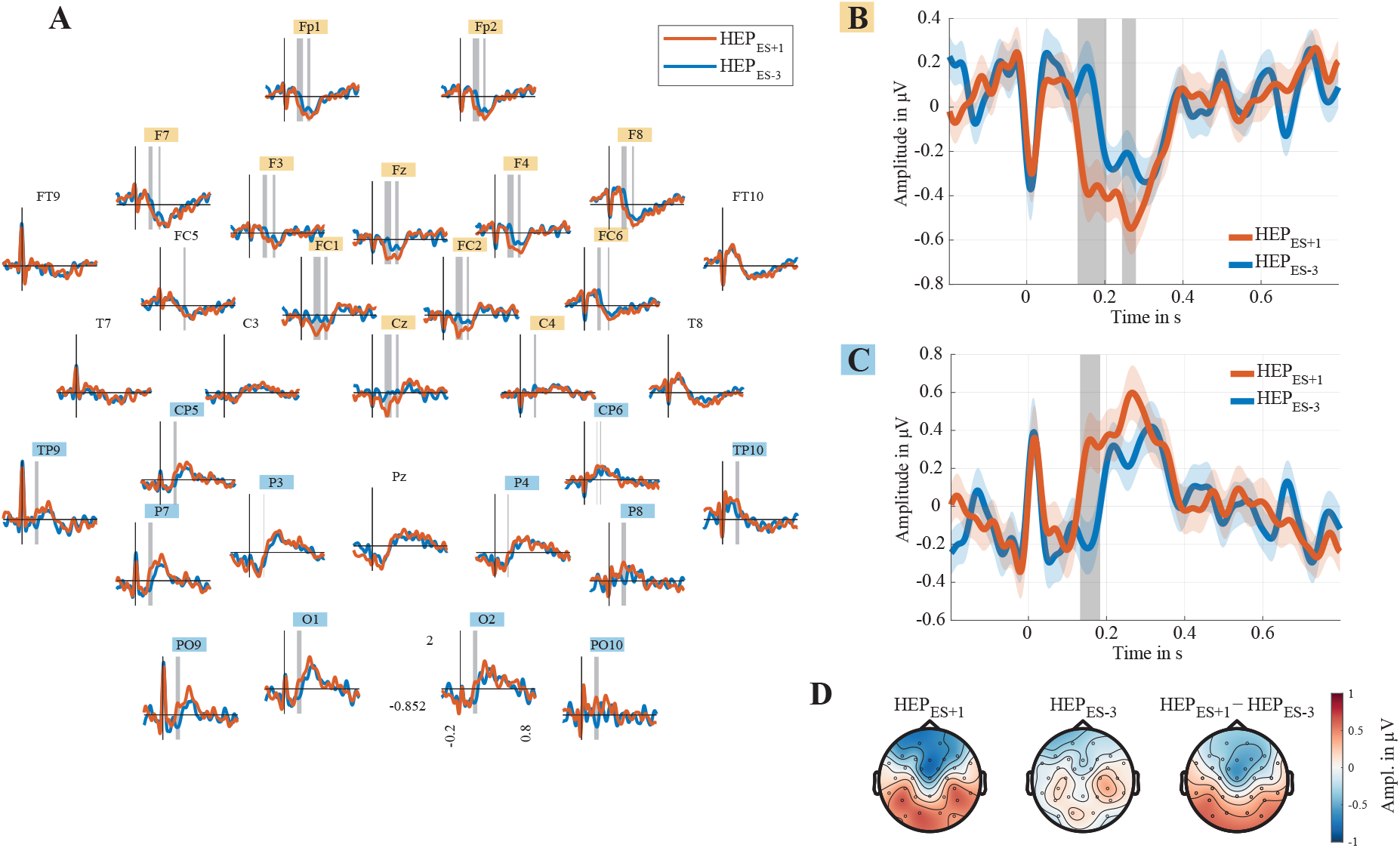
HEP of ES+1 and reference condition (ES-3). Grand average over all participants. **(A)** HEPs in each EEG channel. **(B)** Negative Cluster averaged over central-frontal electrodes (Monte Carlo p = 0.002) between the ES+1 and reference condition. Electrodes used for averaging are highlighted in orange. The grey-shaded area highlights the time window of the found cluster. The coloured-shaded regions around the lines mark the ±1 SEM. **(C)** Positive Cluster averaged over central-frontal electrodes (Monte Carlo p = 0.002) between ES+1 and reference condition. Electrodes used for averaging are highlighted in blue. The grey-shaded area highlights the time window of the found cluster. The coloured-shaded regions around the lines mark the± 1 SEM. **(D)** Topographies of HEPs in ES+1, ES-3 and differences between conditions between 130ms to 200ms after R-peaks.

Both types of ESs were then tested again separately.

#### SVES+1

Running the same statistics revealed a significant difference between SVES+1 and the reference condition (SVES-3) (*p* > 0.05), manifested in a positive and negative cluster from 120 ms to 180 ms. The negative cluster was identified over the central-occipital electrodes (Fp1, Fp2, F3, F4, F7, F8, Fz, FC1, FC2, FC5, FC6, Cz; Monte Carlo *p* = 0.0002), whereas the positive cluster was observed over central-frontal electrodes (T8, FT10, CP5, CP6, TP9, TP10, P4, P7, P8, O1, O2, PO9, PO10; Monte Carlo *p* = 0.008).

#### VES+1

A similar effect was found when comparing VES+1 with the reference condition (VES-3). The cluster-based permutation test showed a significant difference (*p* < 0.05) between the conditions. A negative and a positive cluster were detected in the interval from 140 ms to 210 ms. The negative cluster appeared in the central-occipital electrode region (Fp1, Fp2, F3, F4, F8, Fz, FC1, FC2, FC6, C4, T8, Cz; Monte Carlo *p* = 0.0002) and the positive cluster was located in the central-frontal region (T7, CP5, TP9, P3, P7, P8, O1, O2, PO9, PO10; Monte Carlo *p* = 0.0002).

Because similar temporal and topographic effects were observed for both the VES+1 and SVES+1 conditions, the extent of their similarity was determined. For this purpose, the cluster-based permutation test was performed over the entire period between VES+1 and SVES+1. No significant difference was found between the two ES-groups (*p* > 0.05, Supplementary Figure 18). Therefore, to increase the data set and improve the SNR, they were combined into a single condition (ES+1) for further steps.

### 3.2 Multiverse HEP Analysis

To test the robustness of the results to variations in the pre-processing parameters, different pipelines were used (Figure 4). The same HEP analysis was performed on EEG data that had not been corrected by ICA. Furthermore, the N beats were used as the reference condition and the analysis was performed both without any baseline and with a baseline set immediately before each beat (−150 ms to 50 ms). For all these variations, the VES, ES+1, SVES+1, VES+1 remained significantly different from the reference condition (*p* < 0.05) with similar cluster characteristics (Supplementary Table 3) and thus confirmed the results. Furthermore, slow fluctuations do not significantly affect the HEP in the ES time window because the results remained unchanged when changing the reference condition to N.

**Figure 4:**
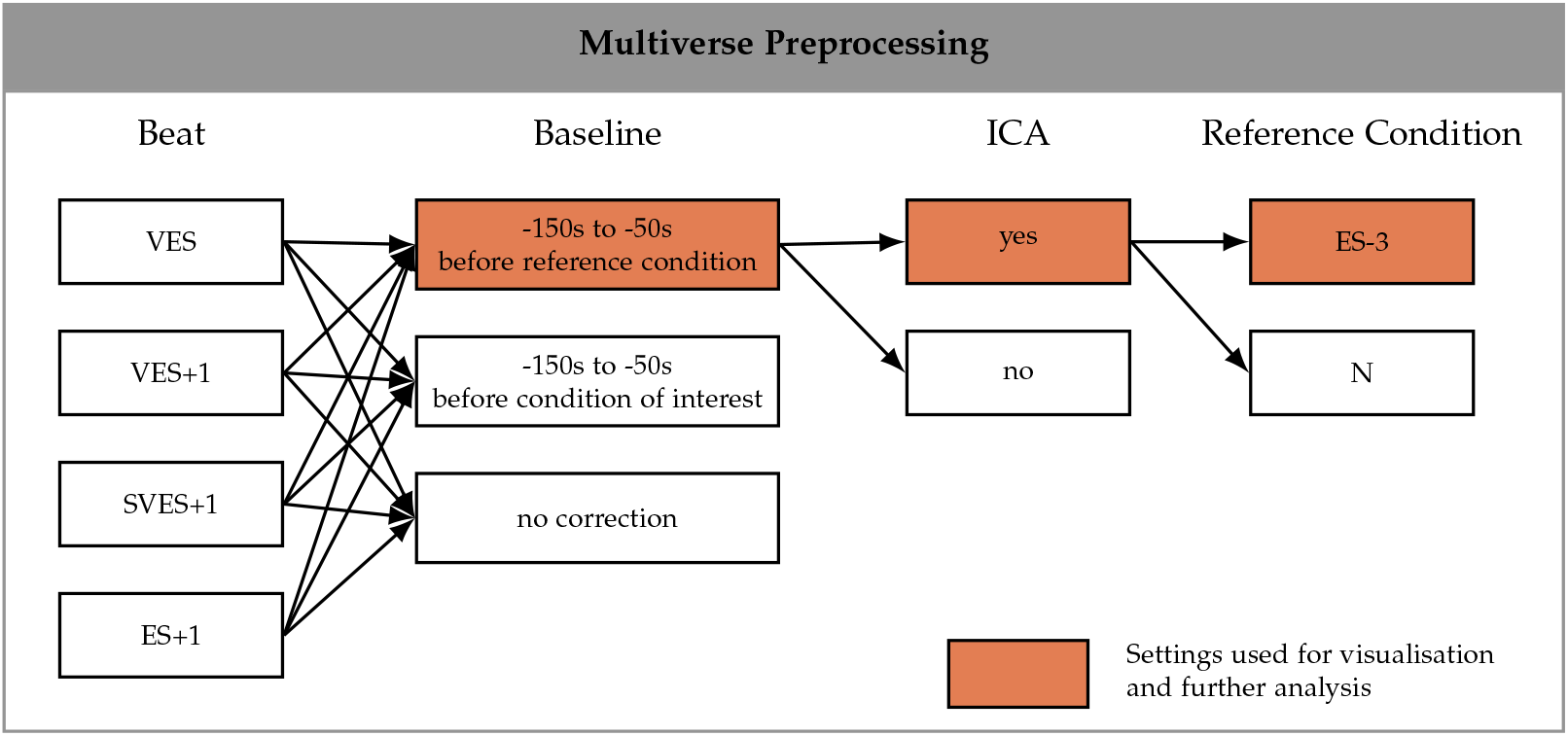
Paths of the multiverse preprocessing pipeline. The settings used for the visualisation and further analysis are highlighted in orange.

### 3.3 Source Space Analysis

The source reconstruction was applied to the averaged HEP within the identified temporal-clusters for all participants separately. Afterwards the sources of the conditions (VES, ES+1) were compared with the sources of the reference condition (ES-3). To obtain robust results localising primary differences in the HEP amplitude between the conditions, source space analysis was performed using different regularisation parameters and aggregation methods average (AVG) and average with signflip (AVG-SF) (Figure 5).

**Figure 5:**
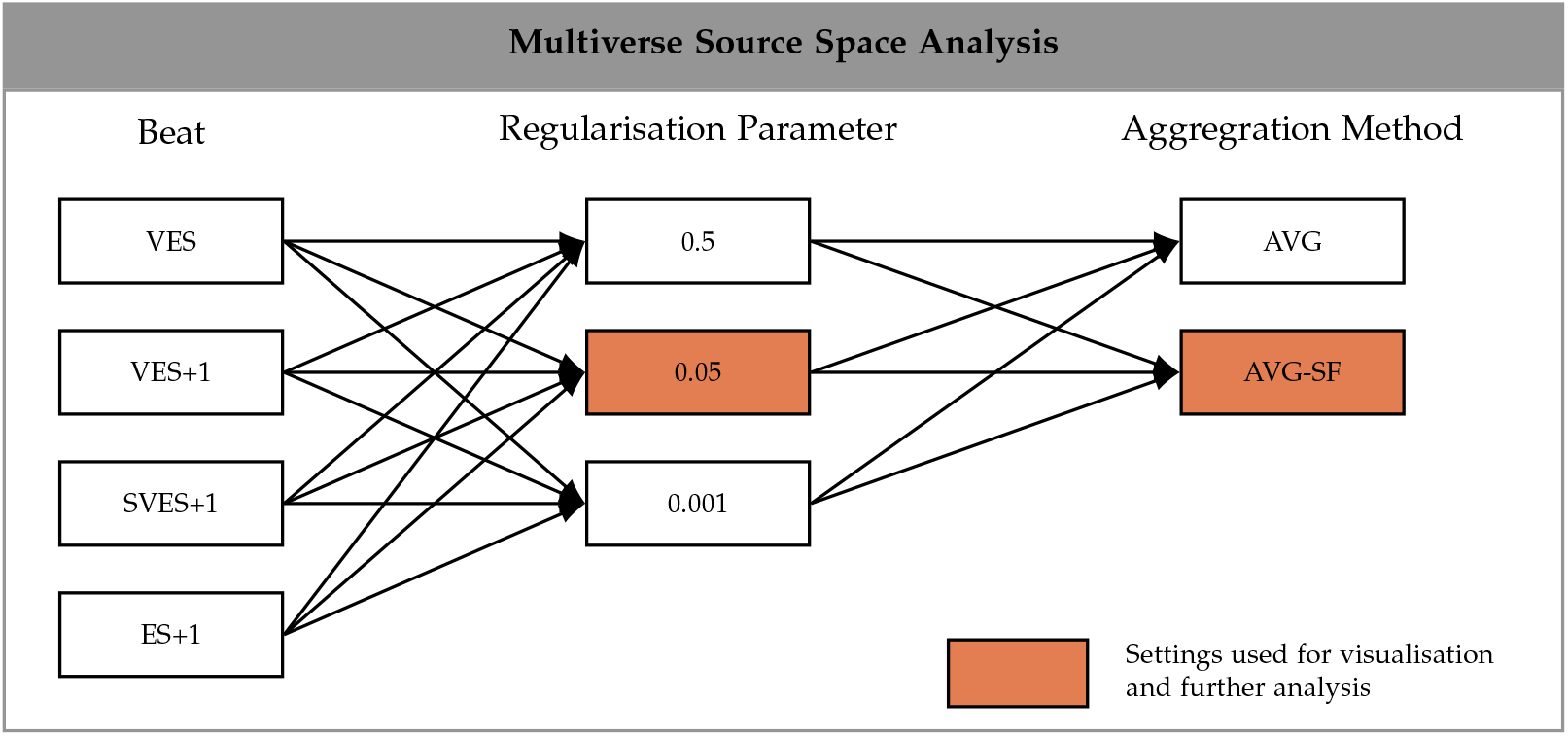
Paths of the multiverse source analysis pipeline. The settings used for the visualisation and further analysis are highlighted in orange.

**Figure 6:**
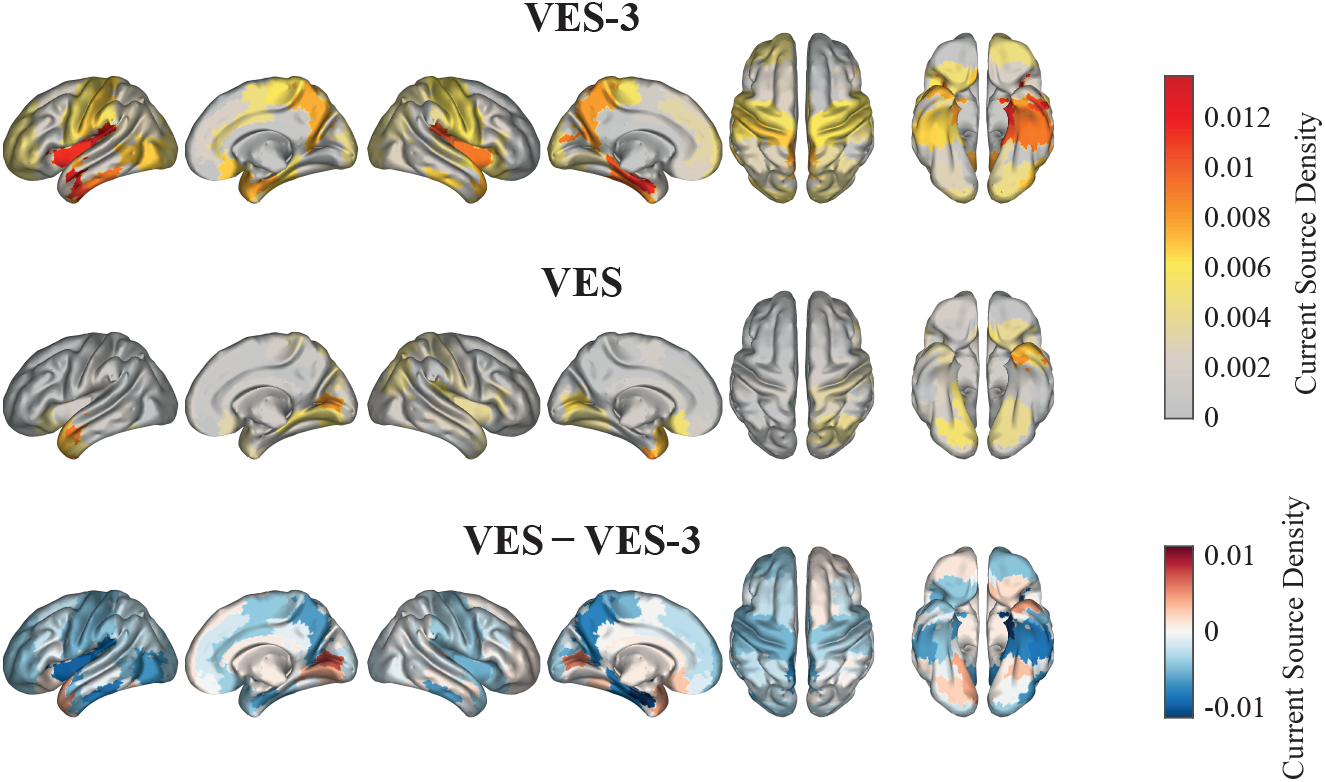
Source Space of HEP separately for VES, VES-3 and difference between conditions in the 220 ms to 350 ms time window.

**Figure 7:**
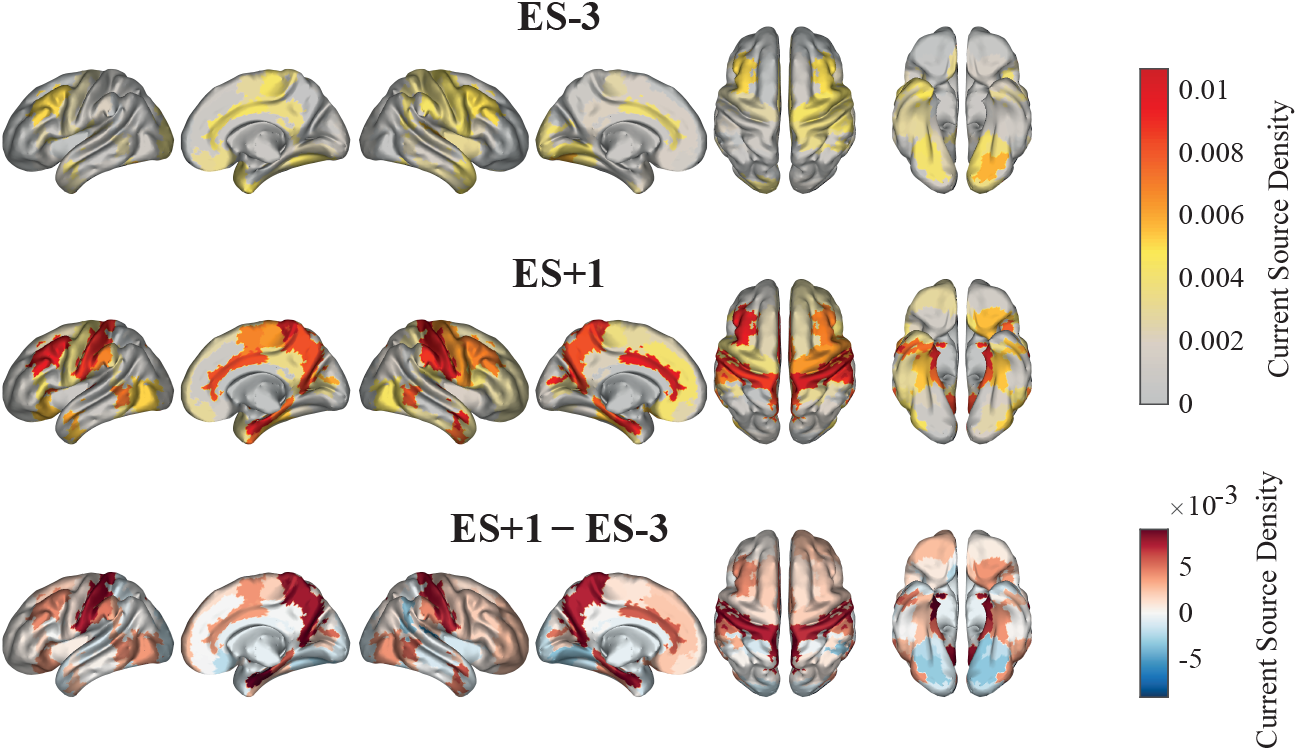
Source Space of HEP separately for ES+1, ES-3 and difference between conditions in the 130 ms to 200 ms time window.

**Figure 8:**
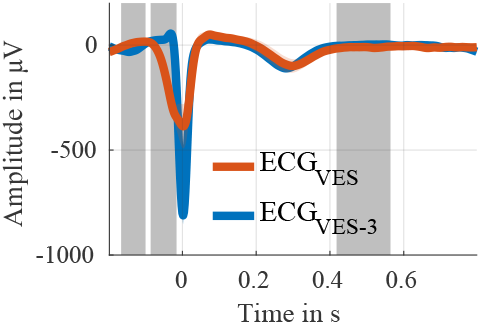
Clusters in ECG between VES and reference condition (Monte Carlo p < 0.0144). The grey-shaded areas highlight the time window of the found clusters. The coloured-shaded areas around the lines mark the ±1 SEM.

**Figure 9:**
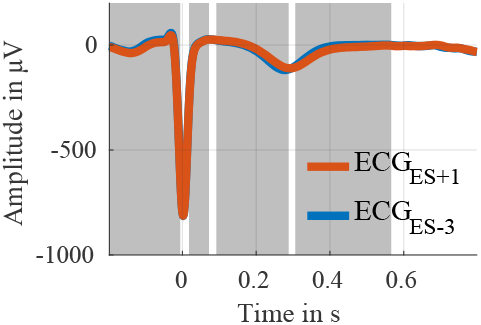
Clusters in ECG between ES+1 and reference condition (Monte Carlo p < 0.023). The grey-shaded areas highlight the time window of the found clusters. The coloured-shaded areas around the lines mark the ±1 SEM.

#### VES

The primary difference in the HEP amplitude between ES/VES and the reference condition across all parameter settings was found in the left insula (*p*_FDR_ < 0.0324, Supplementary Table 5).

#### ES+1

When comparing the HEP amplitude between the ES+1 and ES-3 conditions, significant differences were consistently observed in the left frontal orbital cortex and in both the left and right cingulate gyrus for all parameters (*p*_FDR_ < 0.03, Supplementary Table 4). When both types of ESs were considered separately, the effect remained, except for a few parameter combinations in the SVES+1 condition.

We performed further analyses to determine whether the neuronal response to the ES+1 is a premature and more pronounced HEP response with typical HEP sources or rather a neuronal response showing a distinct source configuration. Therefore, we conducted the same analysis as described above, but compared the ES+1 sources in the time window of 130 ms to 200 ms in multiple comparisons with sources across time windows (130 ms to 200 ms, 200 ms to 270 ms, 270 ms to 340 ms, 340 ms to 410 ms, 410 ms to 480 ms, 480 ms to 550 ms, 550 ms to 620 ms) of the HEP in the ES-3. For this analysis we conducted a one-sided dependent *t*-test for each ROI, to see which sources were significantly more active in the ES+1 condition. We broadened our analysis to include all ROIs rather than limiting it to the predefined HEP ROIs. Across all comparisons, the left and right ACC were significantly more active in the ES+1 condition (*p*_FDR_ < 0.03, Supplementary Figure 19).

### 3.4 ECG Differences

The cluster-based permutation test was applied to the ECG channel under the same conditions to test how potential differences in CFAs might have affected the EEG results.

#### VES

The significant difference (*p* < 0.05) between VES and VES-3 was most pronounced in the time window before the R-peak with a positive cluster from −120 ms to −100 ms (Monte Carlo *p* < 0.0144), a negative cluster from −90 ms to −10 ms (Monte Carlo *p* < 0.0096), and a negative cluster after the T-wave in the time window from 410 ms to 560 ms (Monte Carlo *p* < 0.0012).

#### ES+1

The comparison of the ECG signal between the ES+1 condition and the reference condition (ES-3) showed a significant difference (*p* < 0.05) characterised by three negative clusters (all Monte Carlo *p* < 0.023) and one positive cluster between 100 ms and 290 ms (Monte Carlo *p* = 0.0002). Notably, the T-wave in the ES+1 condition occurs later and with reduced amplitude, in contrast to the results from the EEG analyses. Importantly, this speaks against the differences in ECG as a sole source of the observed effects in the HEP, as the CFAs of the T-wave can not explain the observed differences in the HEP.

### 3.5 Controlling for CFA: Correlation Analysis

Upon identifying significant differences in the ECG data between conditions ES+1 and ES-3, we further conducted a correlation analysis between ECG and EEG time courses to evaluate the potential impact of the differences in the ECG on the observed differences in the HEP. This step was crucial to ensure that the variations detected in EEG signals were not merely reflections of cardiac artefacts but genuine neural activity differences.

#### VES

ECG differences did not significantly correlate with EEG differences over the entire VES time window (Supplementary Figure 20).

Analysing the time window in which main differences between conditions were found (220 ms to 350 ms), the EEG difference at electrode FT9 correlates significantly (*ρ* = −0.49; *p*_FDR_ < 0.05) with the ECG difference of the VES conditions (Supplementary Figure 21). No significant correlation was found for all other electrodes.

This suggests that the effects observed in the EEG for the VES condition are not mainly due to variations in the ECG.

#### ES+1

Performing the same analyses on the ES+1 condition, no significant correlation was found either for the analysis of the entire time window or for the selected averaged time window from 130 ms to 200 ms (*p*_FDR_ > 0.05, Supplementary Figures 22 and 23).

This also implies that the changes observed in the EEG during the ES+1 condition are not primarily due to differences in the ECG.

### 3.6 Control Analyses

To ensure that neither the projection of the T-wave in the ECG signal nor the IBI influenced the HEP amplitude, we matched within-patient epochs in the ES+1 condition to epochs in the N or ES-3 conditions. This matching process involved solving a linear assignment problem with a cost function that allows epochs to remain unmatched, depending on the predefined costs associated with leaving an epoch unmatched. The previously described cluster-based permutation test was then reapplied to the matched epochs on the entire time window. The results of the matching are shown in Supplementary Tables 7 and 6. We only applied the matching for the ES+1 condition, as the impact of the projection of the T-wave on the HEP in the VES condition was already tested in Section 3.1. Furthermore, the matching of the IBI was not used for the VES condition. Neither the IBI from the beat before the ES to the ES nor the IBI from the ES to ES+1 were matchable. This was due to the significant alterations caused by the very short IBI before and the prolonged IBI after the VES.

#### Matching IBI

We matched epochs based on their IBI to investigate if the difference in ES+1 is independent of the IBI. The significant difference in the EEG signal persisted after the matching. We found a negative cluster over central-frontal electrodes (Fp1, Fp2, F3, F4, F7, F8, Fz, FC1, FC2, FC5, FC6, Cz) in the time window of 160 ms to 270 ms (*p* = 0.0002). Simultaneously, there is a positive cluster over central-occipital regions (*p* = 0.0012; CP5, TP10, P3, P4, P7, P8, Pz, O1, O2, PO9, PO10; Supplementary Figure 24).

#### Matching T-Wave Amplitude

Further, we matched epochs based on their T-wave amplitude to control for possible remaining CFA and stroke-volume artefacts. The two conditions remained significantly different. We found one positive (*p* = 0.0004; CP5, CP6, TP9, TP10, P3, P4, P7, P8, O1, O2, PO9, PO10) and one negative cluster (*p* = 0.0002; Fp1, Fp2, F3, F4, F7, F8, Fz, FC1, FC2, FC5, FC6, C4, Cz) approximately between 130 ms to 290 ms (Supplementary Figure 25).

## 4 Discussion

We discovered a significant impact of extrasystoles on heart-brain interaction, as evidenced by altered HEPs during ventricular extrasystolic and both supraventricular and ventricular postextrasystolic beats. More specifically, we found an attenuated neural response to the VES originating in the insular, while the neural response to the SVES showed no significant change. Furthermore, we observed that the neural response to the postextrasystolic beat ES+1 was stronger and earlier than the HEP in the reference condition, indicating a distinct alteration in heart-brain communication following an extrasystolic event. We suggest that our results at least in part can be interpreted within a predictive-coding framework where stronger neural responses are produced by the extrasystolic interruptions of regular rhythmic cardiac cycle.

### 4.1 VES

The HEP was diminished in response to VES, with the most profound attenuation located bilaterally in the insular cortex. The insular cortex is involved in processing interoceptive information, including the perception of the heartbeat [1].

Kumral et al. found a similar attenuation in the HEP in the right insular cortex among patients with atrial fibrillation. They argue that this reflects an altered heart-brain interaction, resulting in an attenuated interoception, which in turn could explain why atrial fibrillation is often reported as asymptomatic or silent. [13] In the context of our study, this finding may help to understand why patients frequently do not experience symptoms during the extrasystole itself.

Furthermore, the baroreceptorreflex, a critical neurophysiological pathway in the heart-brain axis, is attenuated in patients with atrial fibrillation [29] and during VES due to altered hemodynamics [30]. Typically, baroreceptors regulate blood pressure by detecting changes in the stretching of the arterial wall during systole. [1] SVES are associated with less abnormal hemodynamics, potentially leading to a more normal baroreceptorreflex [31, 8, 30]. Accordingly, our findings did not reveal a diminished HEP response in these extrasystoles.

When discussing the changes in the amplitude of HEPs, it is also important to consider cardiac parameters. Specifically, the reduction in stroke volume associated with lower atrial pressure likely leads to reduced activity in the cardiovascular sensory feedback mechanisms, such as baroreceptor activity, somatosensation, and vagal sensory neuron [32] responses, potentially resulting in a decreased HEP amplitude. However, Buot et al. found no positive correlation between HEP amplitude and stroke volume when ICA correction was used. Even though we applied ICA correction, this was based on sinus beats, which may not altogether remove CFAs caused by VES. This suggests that the potentially artifactual correlation between stroke volume and HEP amplitude might still be present in VES.

Another potential artefact affecting the HEP response to VES is the CFA. Especially changes in the T-wave in the ECG can be misinterpreted as a neural response in the EEG due to the late latency of the T-wave. However, due to the polymorphic nature of VES, the CFA within the time frame of diminished HEP amplitude displays both positive and negative polarities. We addressed this by reanalysing the data separately for groups of participants categorised by the polarity of their ECG signals. The results revealed a significant reduction in HEP amplitude in both subsets, indicating that CFAs do not account for the diminished HEP amplitude. Moreover, the polarity of the HEP responses remained the same despite the inverse in polarity of T-wave in ECG recordings.

### 4.2 ES+1

In the context of the postextrasystolic beat of VES and SVES, the HEP amplitude was significantly larger in the time window of 130 ms to 200 ms, originating from sources in the left orbitofrontal cortex and the ACC, areas known for their involvement in emotional processing and autonomic control [33].

The concept of predictive coding offers a valuable framework for interpreting these findings. The HEP is thought to be processed through these predictive coding mechanisms, with the HEP amplitude reflecting a prediction error [34, 4, 35]. According to the framework, the brain continuously predicts incoming sensory information from exteroceptive and interoceptive signals [33] based on past experiences. When these predictions are not met, it results in a prediction error, prompting the brain to update its model of the world. In interoceptive processing, the brain anticipates the heartbeat’s regular strength, rhythm, and variability. This implies that the brain predicts the heartbeat given the history of the previous heart beats. [34] A bottom-up deviation in heart functioning, such as a disrupted sinus rhythm by an ES, does not match the top-down prediction, generating a prediction error that necessitates updating the model. ESs often lead to a stronger sensation in the subsequent heartbeat [9], potentially further highlighting this mismatch between the prediction and the actual sensation.

Essential sources for processing these prediction signals are the anterior insular cortex, ACC, and orbitofrontal cortex. Medford and Critchley state that the ACC and anterior insular coordinate responses to internal and external events. [36] Specifically in interoception, the ACC is involved in error detection and integrating emotional predictions [33]. Our analysis indicated that especially the ACC is the underlying destinct source of the earlier and stronger HEP, further suggesting that the HEP reflects a prediction error. Yoshimoto et al. also found that the ACC may serve as a top-down centre for heart rate regulation [37], which is crucial especially after an extrasystole [38].

Notably, the identified time window of 130 ms to 200 ms is earlier than the typically reported HEP window of 200 ms to 600 ms [1]. It aligns more closely with the time window associated with MMN, typically found between 100 ms and 250 ms. MMN represents a neural marker in the predictive coding framework for detecting deviations from expected sensory patterns. [39] The mentioned conceptual and temporal alignments raise the possibility of a similar underlying mechanism.

Previous studies have demonstrated a prediction error in HEPs related to cross-modal predictive processing. For example, Pfeiffer et al. found a neural response to unexpected omissions in auditory sequences synchronised with the heartbeat. They observed a slightly earlier and larger HEP amplitude over the posterior-central scalp regions, suggesting that the brain’s capacity for regularity processing extends to interoceptive signals. [4] Furthermore, Banellis and Cruse observed similar effects modulated by attention and perceived synchronicity within an early time window of the HEP (27 ms to 230 ms) overlapping with the time window found in this study [5]. Our research aligns with these findings, as the disturbed rhythm due to the occurrence of an extrasystole represents an unexpected event, possibly eliciting a surprise response. Consequently, our study goes beyond the cross-modal predictive mechanism and demonstrates a prediction error in HEP in the same interoceptive modality—the heartbeat itself.

The exact nature of the predicted “heartbeat” event remains unclear. In auditory MMN studies, where the MMN is detected in response to prematurely presented stimuli, the MMN generally occurs at the moment of the unexpected stimulus itself rather than afterwards. [3] However, other types of deviants, like frequency, strength or duration, can also elicit MMN-like responses in a variety of modalities like visual and somatosensory. This leaves uncertainty about whether the MMN detected in our study is linked to the surprise of the long compensatory pause after the extrasystole or even the increased stroke volume of the postextrasystolic heartbeat. Further research is needed to clarify the specific predictions made by the brain regarding heartbeats and how it processes deviations from these predictions.

MMNs are triggered pre-attentively, meaning without consciousness [39]. This aligns with the commonly asymptomatic nature of extrasystoles [9]. These parallels further support the hypothesis that the HEP for the postextrasystolic beat can be categorized within predictive coding and embodies a prediction error similar to an MMN. Ehlers et al. showed that the heartbeat perception of patients with extrasystoles is not significantly different from that of the control group [40]. However, they only investigated normal heartbeat perception, not the perception of extrasystoles, which might differ. Further research is needed to assess the interoceptive perception of extrasystoles and its relation to the HEP.

If perceived, anxiety is a common symptom of extrasystoles [40, 41]. It has been earlier hypothesised that anxiety may arise due to the mismatch between predicted and actual interoceptive stimuli [42]. Garfinkel et al. have shown that an interoceptive mismatch is associated with heightened anxiety symptomatology [43], which further supports the idea that the anxiety experienced during extrasystoles could be indicative of an interoceptive mismatch mechanism. Given that a regular heartbeat is crucial for survival, deviations from this regularity, beyond normal heart rate variability, can have fatal consequences. A signal, such as a prediction error, could alert the individual to potential internal threats, such as an abnormally prolonged pause in the rhythm. This mechanism could explain the heightened anxiety experienced during such events.

While we corrected and controlled for heartbeat artefacts, residual effects may have influenced the reported findings. The postextrasystolic beat differs in cardiac parameters, including a significant difference in the ECG and likely in the CFA. However, we observed an earlier and stronger EEG amplitude compared to a normal HEP, but the T-wave, which falls in the critical HEP time window, appears later, known as a QT-prolongation. [44, 45] Hence, it is unlikely that a later CFA introduces these earlier differences in the EEG. Moreover, we found that the differences in the ECG do not correlate with those in the EEG, and the effects observed are independent of the T-wave amplitude and IBIs, indicating that our findings are not simply artefacts.

However, in our study, we used the IBI from ES+1 to ES+2 to match the IBIs of ES+1 with those of regular sinus beats. This strategy was adopted to circumvent the challenge of matching the longer IBI interval from ES to ES+1, known as the compensatory pause, with normal sinus beats due to difficulty finding adequate matches. This approach mitigates the problem of insufficient matching partners but does not consider potential artefacts related to stroke volume. The compensatory pause length in extrasystoles is positively correlated with the stroke volume of the subsequent post-extrasystolic beat (ES+1) [27, 46], an aspect our matching process does not adjust for due to the scarcity of suitable matches. Consequently, while our method effectively controls for CFAs that might vary with the IBIs of the current beat, it does not address variations in stroke volume. However, we reasoned that if a stroke volume would be a sole reason for larger HEPs to ES+1 heart beats, then we should see an increase in the amplitude of source activity typically associated with HEPs. However, the HEP sources to ES+1 beats were generated in different cortical, especially the ACC, known as central for processing prediction errors. This indicates that a mere scaling of normal HEPs does not represent a neural processing relating to ES+1 beats.

A further, possibly complementary, explanation for the increased HEP amplitude might involve the increased firing of baroreceptors in response to the increased stroke volume. However, the information from the baroreceptors’ cardiac afferents after a sinus beat is centrally processed 400 ms to 800 ms after the ECG R-peak [1]. This time frame lies remarkably after the observed effect. Furthermore, the increased baroreceptor activity does not account for the earlier timing of neural response. Considering the cardiac stroke volume as a potential artefact, it correlates with the HEP amplitude at around 500 ms post-R-peak [1]. However, the observed effect in the HEP occurs much earlier when stroke volume and should not influence the HEP. Consequently, this effect might not be solely an artefact of stroke volume but a genuine neural response. Still, further investigation into the mechanisms underlying the observed HEP amplitude increase and its temporal dynamics by recording baroreceptor firing and the pulse wave would provide valuable insights.

The observed effects might also result from the increased contractility of the postextrasystolic beat [47], leading to increased/earlier firing of cardiac neurons or somatosensory pathways, which could provide a faster response than the slower baroreceptors. One direct neural pathway involves cardiac neurons in the heart wall that detect chemical and mechanical changes in the atria and ventricles. These neurons have fast conduction velocities, observable in the HEP, within relatively early time windows (e.g., shortly after R-waves, <100 ms). Similarly, somatosensory pathways, including mechanoreceptors in the skin and muscles, could convey information about cardiac interoception, providing a more immediate neural response than slower baroreceptor-mediated pathways. [48] These alternative pathways align with the earlier timing of the neural response observed in the study. However, the findings of the source analysis indicate that the effect observed in the postextrasystolic beat is not merely a stronger HEP due to fast pathways. The activity of sources is closely linked to the underlying neural mechanisms occurring during the corresponding phase of the HEP. If the early difference in HEP were solely due to a stronger and earlier cardiac signal, we would only observe a stronger typical HEP source during this earlier time window. However, we identified the ACC being exclusively active during this early time window of the postextrasystolic beat. The ACC was not active in the same early time window of the reference condition, nor was it active at the time of the highest amplitude or any other time window during the HEP of the reference condition. This indicates that the observed effect is not merely a stronger and earlier HEP response but rather an independent mechanism. However, studies involving direct recordings from cardiac neurons and somatosensory pathways during postextrasystolic beats could provide valuable insights into their roles in modulating HEPs.

Another yet-to-be-explored indicator for MMN-like processes could be the correlation between the strength of the perception of the post-extrasystolic beat and the magnitude of the change in the HEP. Therefore, it would be advisable to analyse the interoceptive perception of the postextrasystolic beat using participant feedback about the timing and intensity of the perception. Such a correlation would indicate that stronger perceptions of the post-extrasystolic beat are associated with larger changes in the HEP, mirroring the way MMN reflects the brain’s response to deviations from expected exteroceptive patterns. By understanding this correlation, we could better comprehend how the brain processes unexpected interoceptive events and their impact on emotional states in a behavioural way.

In summary, our findings align with the predictive coding framework, indicating that naturally occurring interoceptive stimuli, such as a disruption in heart rhythm due to extrasystoles, induce prediction errors in the ACC, thereby requiring updates of the brain’s internal model and deepening our comprehension of interoceptive processing

