## Supplementary for "Neural Representations of Extrasystoles: A Predictive Coding Perspective"

### A Appendix

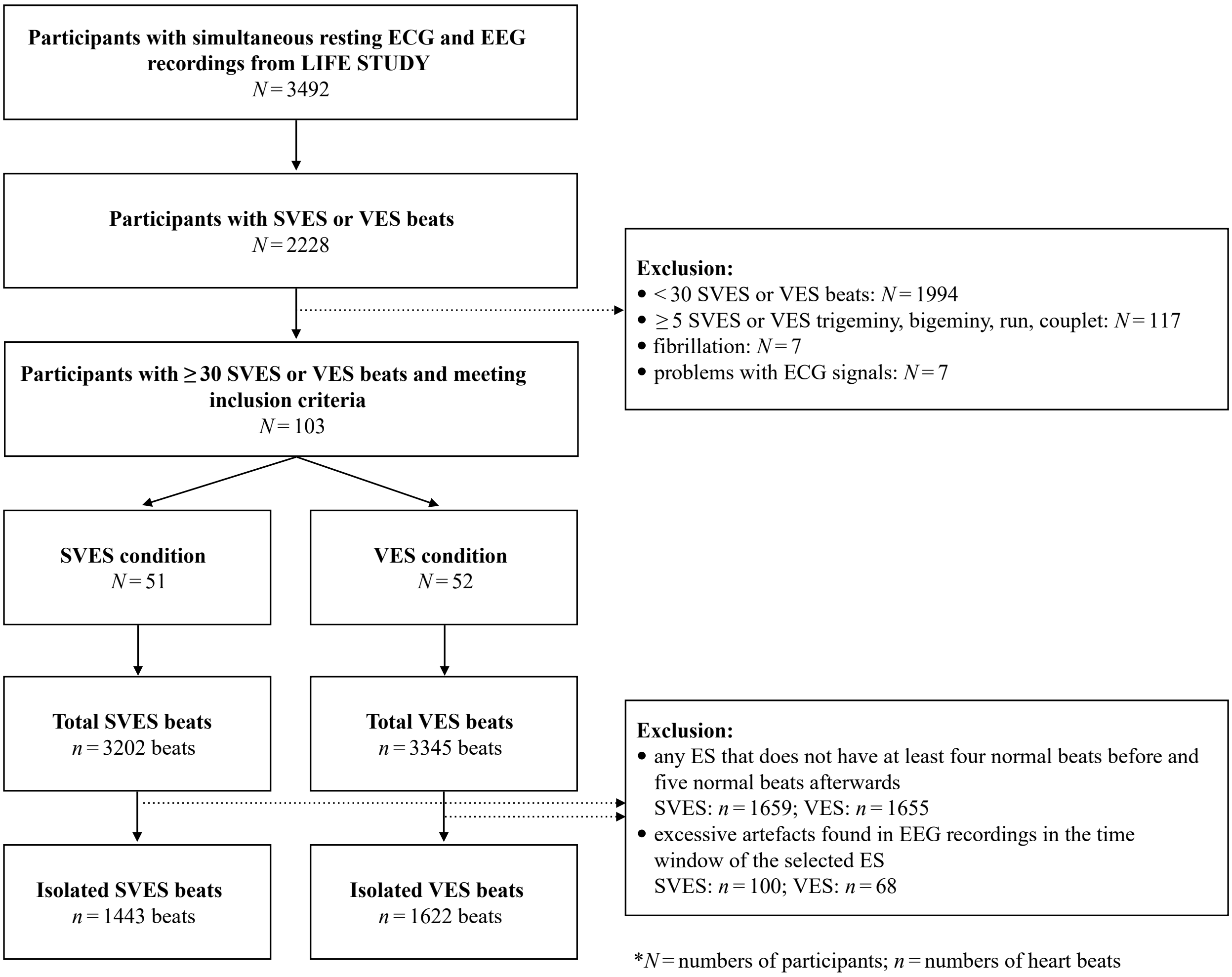


**Figure 10:** *Flow chart of the selection of participants with ES beats.*

**Table 1:** *IBIs of beats surrounding and including SVES.*

| SVES | IBI in ms | | | | | | | |
|---|---|---|---|---|---|---|---|---|
| | Sinusbeat | SVES-4 | SVES-3 | SVES-2 | SVES-1 | SVES | SVES+1 | $\frac{IBI_{SVES-1}+IBI_{SVES}}{2 \cdot IBI_{Sinusbeat}}$ in % |
| mean | 889.34 | 924.76 | 921.69 | 926.39 | 578.38 | 1114.99 | 950.88 | 0.96 |
| std | 131.36 | 148.51 | 147.25 | 139.97 | 90.12 | 165.82 | 158.42 | 0.06 |
| min | 594.53 | 593.11 | 587.78 | 587.78 | 438.44 | 713.11 | 608.89 | 0.80 |
| max | 1205.76 | 1285.82 | 1218.80 | 1216.62 | 826.00 | 1478.55 | 1442.18 | 1.08 |

**Table 2:** *IBIs of beats surrounding and including VES.*

| VES | IBI in ms | | | | | | | $\frac{IBI_{VES-1}+IBI_{VES}}{2 \cdot IBI_{Sinusbeat}}$ in % |
|---|---|---|---|---|---|---|---|---|
| | **Sinusbeat** | **VES-4** | **VES-3** | **VES-2** | **VES-1** | **VES** | **VES+1** | |
| mean | 895.76 | 938.60 | 938.56 | 941.27 | 574.09 | 1260.00 | 931.10 | 1.02 |
| std | 111.99 | 121.33 | 120.62 | 122.21 | 111.33 | 205.09 | 132.66 | 0.05 |
| min | 591.30 | 597.09 | 598.91 | 596.73 | 449.81 | 682.55 | 593.09 | 0.83 |
| max | 1222.92 | 1242.32 | 1240.63 | 1246.11 | 1037.09 | 1650.35 | 1249.05 | 1.13 |

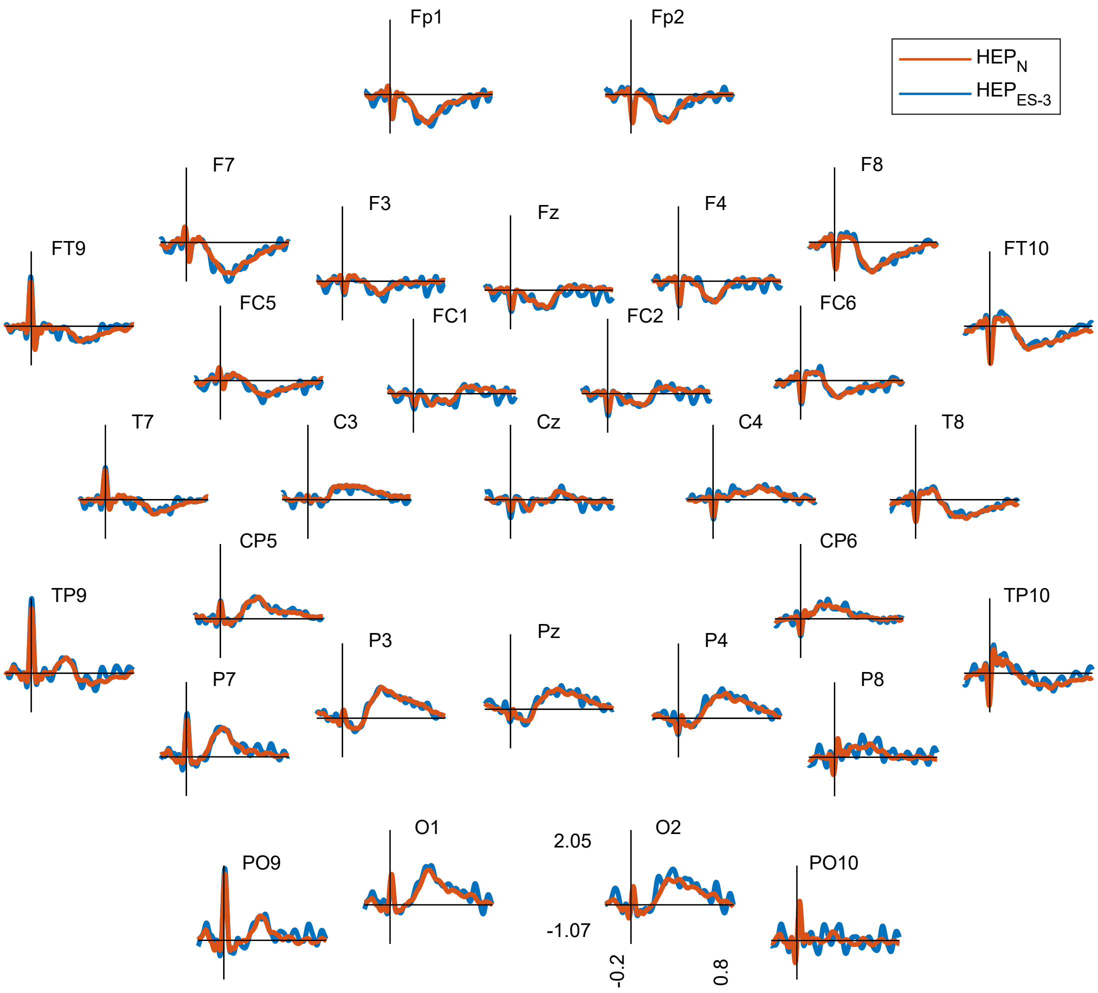


**Figure 11:** *No significant differences ($p > 0.3$) in HEP in the time domain between condition ES-3 and N.*

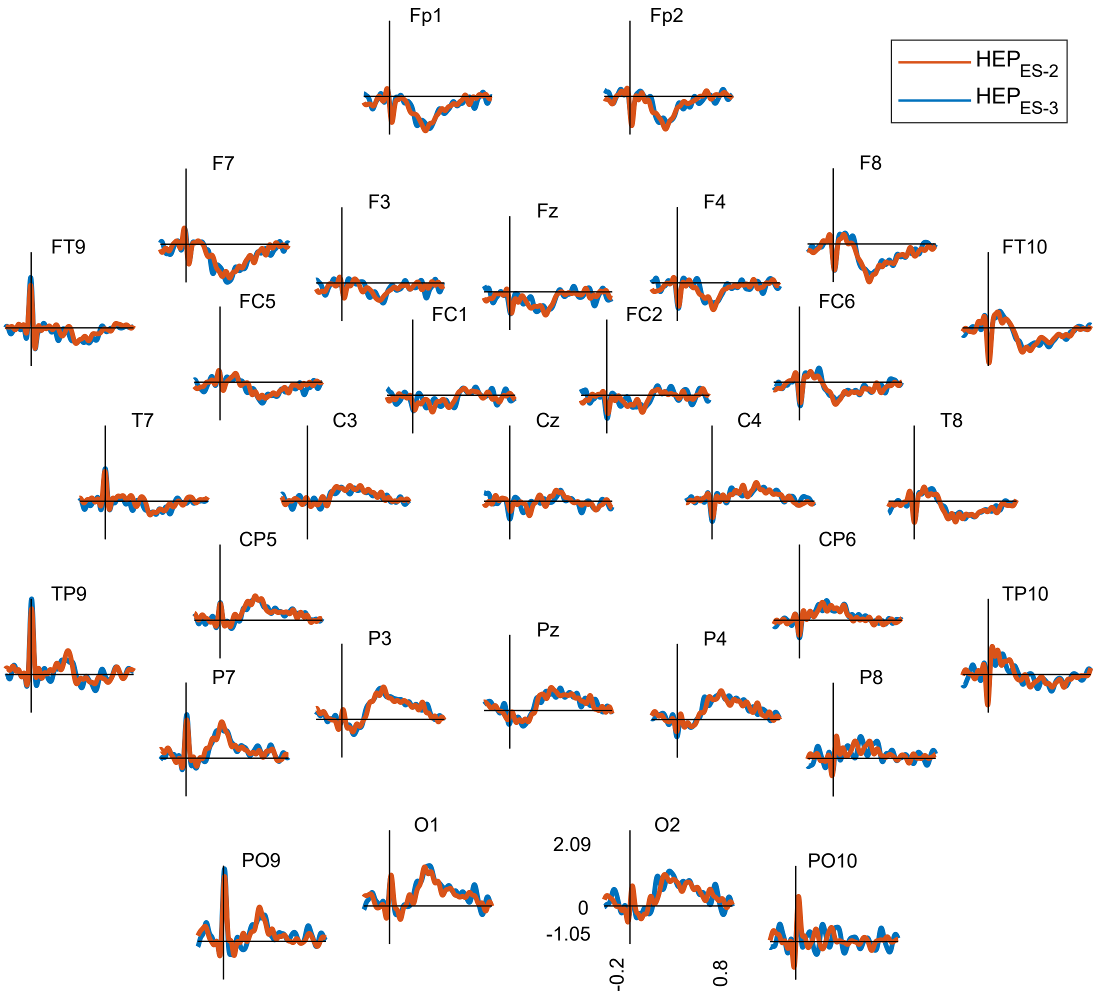


**Figure 12:** *No significant differences ($p > 0.7$) in HEP in the time domain between condition ES-2 and ES-3.*

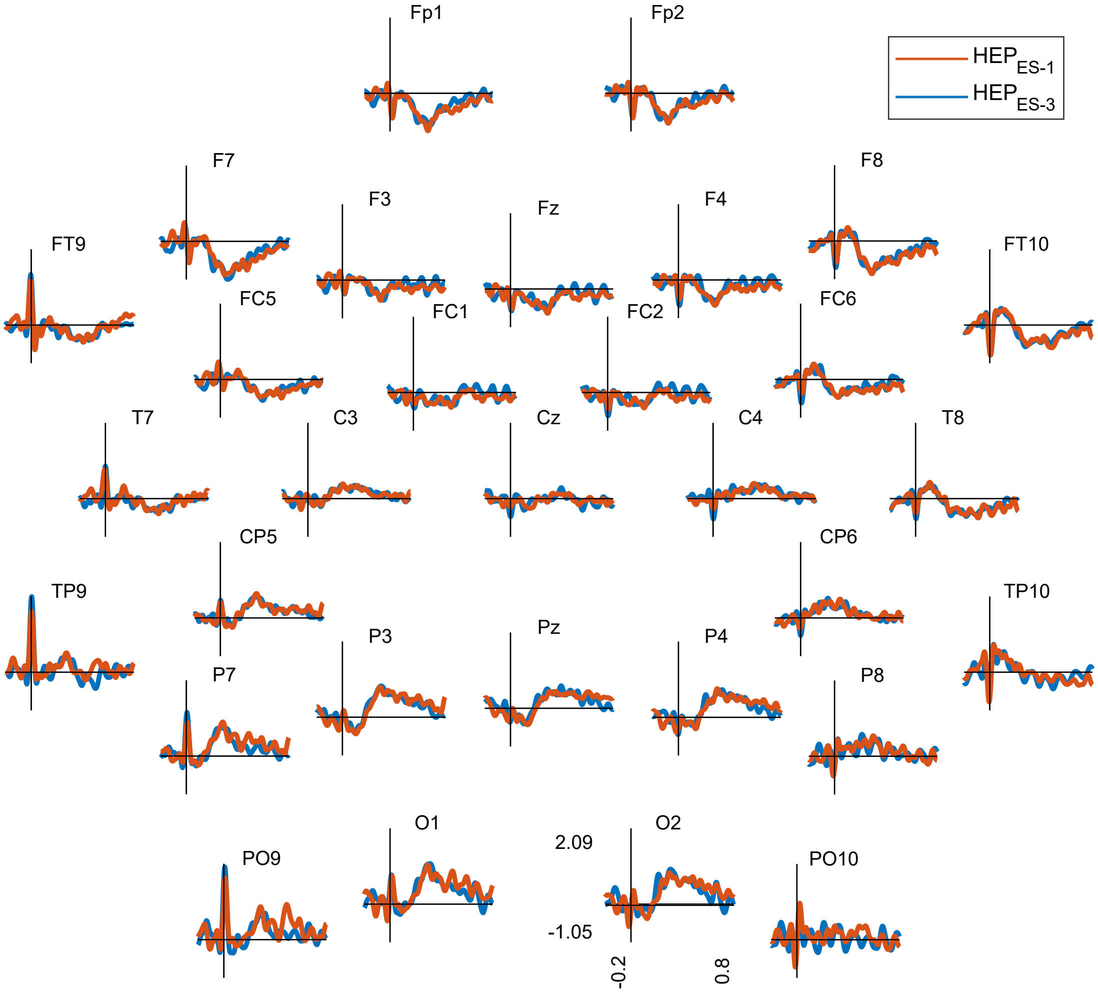


**Figure 13:** *No significant differences ($p > 0.3$) in HEP in the time domain between condition ES-1 and ES-3.*

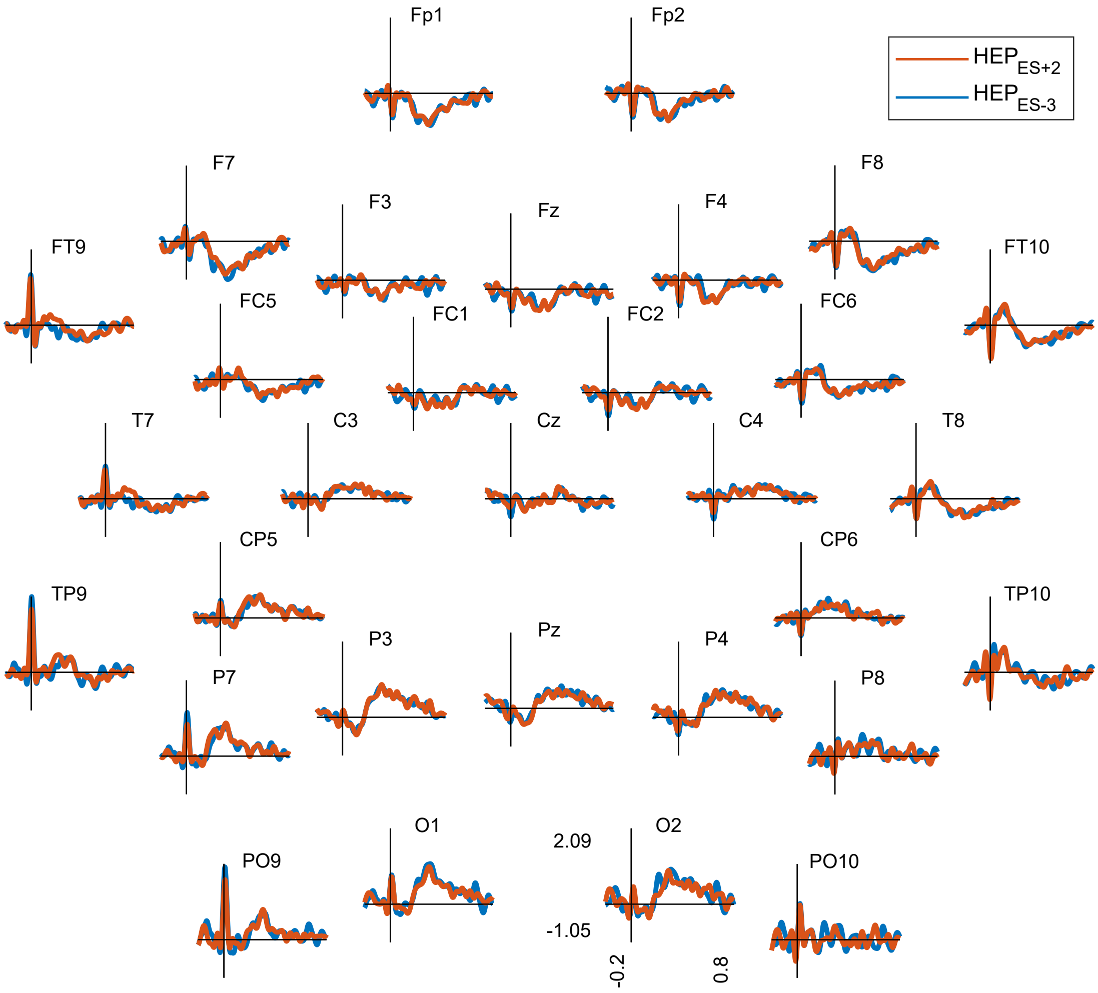


**Figure 14:** *No significant differences ($p > 0.7$) in HEP in the time domain between condition ES+2 and ES-3.*

| | Baseline | Reference Condition | ICA | significant | latency | Cluster electrodes | polarity | sum(t) | Monte Carlo $p$ |
|---|---|---|---|---|---|---|---|---|---|
| VES | −150 ms to −50 ms before reference condition | VES-3 | yes | yes | 224 ms to 332 ms | C3,CP5,CP6,TP9,TP10,P3,P4,P7,P8,O1,O2,PO9,PO10 | negative | -916 | 0.0008 |
| | | | | | 226 ms to 348 ms | Fp1,Fp2,F3,F4,F7,F8,Fz,FC1,FC5,FC6,C3,T7,FT9 | positive | 1348 | 0.0002 |
| VES+1 | | VES-3 | | yes | 138 ms to 204 ms | Fp1,Fp2,F3,F4,F8,Fz,FC1,FC2,FC6,C4,T8,Cz | negative | -1115 | 0.0004 |
| | | | | | 146 ms to 214 ms | T7,CP5,TP9,P3,P7,P8,O1,O2,PO9,PO10 | positive | 806 | 0.0004 |
| SVES+1 | | SVES-3 | | yes | 126 ms to 178 ms | Fp1,Fp2,F3,F4,F7,F8,Fz,FC1,FC2,FC5,FC6,Cz | negative | -760 | 0.0002 |
| | | | | | 128 ms to 180 ms | T8,FT10,CP5,CP6,TP9,TP10,P4,P7,P8,O1,O2,PO9,PO10 | positive | 464 | 0.0013 |
| ES+1 | | ES-3 | | yes | 128 ms to 208 ms | Fp1,Fp2,F3,F4,F7,F8,Fz,FC1,FC2,FC6,C4,Cz | negative | -1454 | 0.0002 |
| | | | | | 132 ms to 186 ms | CP5,CP6,TP9,TP10,P3,P4,P7,P8,O1,O2,PO9,PO10 | positive | 800 | 0.0002 |
| VES | −150 ms to −50 ms before condition of interest | VES-3 | yes | yes | 224 ms to 280 ms | C3, CP5, TP9, P3, P7, O1, PO9 | negative | -417 | 0.0053 |
| | | | | | 236 ms to 308 ms | Fp1, Fp2, F3, F4, F7, F8, Fz, FC1, FC5, FC6, T7 | positive | 549 | 0.0194 |
| VES+1 | | VES-3 | | yes | 136 ms to 196 ms | Fp1, Fp2, F3, F4, F8, Fz, FC1, FC2, FC6, C4, Cz | negative | -831 | 0.0006 |
| | | | | | 144 ms to 212 ms | CP5, TP9, P3, P4, P7, P8, Pz, O1, O2, PO9, PO10 | positive | 720 | 0.0016 |
| SVES+1 | | SVES-3 | | yes | 126 ms to 174 ms | Fp1, Fp2, F3, F4, F7, F8, Fz, FC1, FC2, FC5, FC6, Cz | negative | -641 | 0.0024 |
| | | | | | 126 ms to 172 ms | CP5, TP9, P4, P7, P8, Pz, O1, O2, PO9, PO10 | positive | 418 | 0.0244 |
| ES+1 | | ES-3 | | yes | 130 ms to 196 ms | Fp1, Fp2, F3, F4, F7, F8, Fz, FC1, FC2, FC5, FC6, C4, Cz | negative | -1175 | 0.0002 |
| | | | | | −200 ms to −156 ms | Fp1, Fp2, F3, F4, F7, Fz, FC1, FC2, FC5, C3, Cz | negative | -449 | 0.0236 |
| | | | | | 128 ms to 278 ms | T7, CP5, TP9, TP10, P3, P4, P7, P8, Pz, O1, O2, PO9, PO10 | positive | 1509 | 0.0002 |
| VES | no baseline correction | VES-3 | yes | yes | 224 ms to 332 ms | C3, CP5, CP6, TP9, TP10, P3, P4, P7, P8, O1, O2, PO9, PO10 | negative | -916 | 0.0014 |
| | | | | | 226 ms to 348 ms | Fp1, Fp2, F3, F4, F7, F8, Fz, FC1, FC5, FC6, C3, T7, FT9 | positive | 1348 | 0.0002 |
| VES+1 | | VES-3 | | yes | 138 ms to 204 ms | Fp1, Fp2, F3, F4, F8, Fz, FC1, FC2, FC6, C4, T8, Cz | negative | -1115 | 0.0002 |
| | | | | | 146 ms to 214 ms | T7, CP5, TP9, P3, P7, P8, O1, O2, PO9, PO10 | positive | 806 | 0.0002 |
| SVES+1 | | SVES-3 | | yes | 128 ms to 176 ms | Fp1, Fp2, F3, F4, F7, F8, Fz, FC1, FC2, FC5, FC6, Cz | negative | -683 | 0.0006 |
| | | | | | 134 ms to 174 ms | CP5, CP6, TP9, TP10, P7, P8, O1, O2, PO9, PO10 | positive | 382 | 0.0230 |
| ES+1 | | ES-3 | | yes | 130 ms to 204 ms | Fp1, Fp2, F3, F4, F7, F8, Fz, FC1, FC2, FC6, C4, Cz | negative | -1344 | 0.0002 |
| | | | | | 244 ms to 280 ms | Fp1, Fp2, F3, F4, F7, F8, Fz, FC1, FC2, FC5, FC6, Cz | negative | -447 | 0.0134 |
| | | | | | 132 ms to 184 ms | CP5, CP6, TP9, TP10, P3, P4, P7, P8, O1, O2, PO9, PO10 | positive | 766 | 0.0008 |
| VES | −150 ms to −50 ms before reference condition | VES-3 | no | yes | 228 ms to 374 ms | C4, CP5, CP6, TP10, P3, P4, P7, P8, Pz, O1, O2, PO9, PO10 | negative | -1662 | 0.0002 |
| | | | | | −118 ms to −484 ms | Fp2, F4, F8, FC2, FC6, C4, T8, FT10, CP6, TP10, P8, O2, PO10 | negative | -1311 | 0.0010 |
| | | | | | 146 ms to 484 ms | Fp1, Fp2, F3, F4, F7, Fz, FC1, FC5, T7, FT9, TP9 | positive | 2682 | 0.0002 |
| | | | | | −54 ms to −16 ms | F7, FC5, C3, T7, FT9, CP5, TP9, P3, P7, O1, PO9 | positive | 673 | 0.0002 |
| VES+1 | | VES-3 | | yes | 136 ms to 214 ms | F4, F8, Fz, FC1, FC2, FC6, C4, Cz | negative | -834 | 0.0016 |
| | | | | | −200 ms to −128 ms | F3, FC1, C3, Cz, P3, P4, Pz | negative | -539 | 0.0128 |
| | | | | | −48 ms to 40 ms | F7, FC5, C3, T7, FT9, CP5, TP9, P3, P7, O1, PO9 | negative | -533 | 0.0132 |
| | | | | | 142 ms to 216 ms | FC1, Cz, P3, P4, Pz, O2 | positive | 660 | 0.0056 |
| SVES+1 | | SVES-3 | | yes | −200 ms to −112 ms | FC1, FC2, C3, Cz, P3, P4, P7, Pz, O1, O2, PO9 | negative | -887 | 0.0014 |
| | | | | | 100 ms to 228 ms | F8, FC6, T8, FT10, CP6, TP10, P8, PO10 | positive | 1017 | 0.0020 |
| | | | | | −4 ms to 82 ms | F8, FC6, T8, FT10, CP6, TP10, P8, PO10 | positive | 640 | 0.0092 |
| ES+1 | | ES-3 | | yes | 14 ms to 298 ms | F4, F8, Fz, FC1, FC2, FC6, C3, C4, Cz, P3, P4, Pz, O2 | negative | -2292 | 0.0002 |
| | | | | | −200 ms to −4 ms | FC1, FC2, C3, C4, Cz, CP5, P3, P4, Pz, O1, O2 | negative | -1971 | 0.0004 |
| | | | | | 126 ms to 214 ms | T7, CP5, CP6, TP9, TP10, P7, P8, O1, PO9, PO10 | positive | 910 | 0.0028 |
| VES | −150 ms to −50 ms before reference condition | N | yes | yes | 222 ms to 302 ms | C3, CP5, CP6, TP9, TP10, P3, P4, P7, P8, O1, O2, PO9, PO10 | negative | -799 | 0.0058 |
| | | | | | 240 ms to 306 ms | Fp1, Fp2, F3, F4, F7, F8, Fz, FC1, FC2, FC5, FC6, T7, FT9 | positive | 841 | 0.0048 |
| VES+1 | | N | | yes | 140 ms to 212 ms | Fp1, Fp2, F3, F4, F7, F8, Fz, FC1, FC2, FC6, Cz | negative | -798 | 0.0014 |
| | | | | | 138 ms to 220 ms | CP5, TP9, TP10, P3, P4, P7, P8, Pz, O1, O2, PO9, PO10 | positive | 752 | 0.0014 |
| SVES+1 | | N | | yes | 126 ms to 216 ms | Fp1, Fp2, F3, F4, F8, Fz, FC1, FC2, FC6, Cz | negative | -1037 | 0.0002 |
| | | | | | 42 ms to 112 ms | Fz, FC1, FC2, Cz, P3, P4, Pz | negative | -438 | 0.0104 |
| | | | | | 236 ms to 276 ms | Fp1, Fp2, F3, F4, F8, Fz, FC1, FC2, FC6, Cz | negative | -434 | 0.0110 |
| | | | | | 132 ms to 172 ms | CP5, CP6, TP9, TP10, P3, P7, P8, O1, O2, PO9, PO10 | positive | 415 | 0.0168 |
| ES+1 | | N | | yes | 124 ms to 288 ms | Fp1, Fp2, F3, F4, F7, F8, Fz, FC1, FC2, FC5, FC6, Cz | negative | -2513 | 0.0002 |
| | | | | | 130 ms to 286 ms | CP5, CP6, TP9, TP10, P3, P4, P7, P8, Pz, O1, O2, PO9, PO10 | positive | 1946 | 0.0002 |

**Table 3:** *Multiverse Preprocessing Analysis of HEP in the time domain.*

| regularisation parameter | aggregation method | target condition | reference condition | Left Cingulate Gyrus positiveterior division | Right Cingulate Gyrus positiveterior division | Left Postcentral Gyrus | Right Postcentral Gyrus | Left Frontal Orbital Cortex | Right Frontal Orbital Cortex | Left Insular Cortex | Right Insular Cortex | Left Cingulate Gyrus anterior division | Right Cingulate Gyrus anterior division |
|---|---|---|---|---|---|---|---|---|---|---|---|---|---|
| 0.5 | AVG | ES+1 | ES-3 | 0.3717 | 0.9469 | 0.1305 | 0.0796 | 0.0000 | 0.0000 | 0.6906 | 0.0796 | 0.0000 | 0.0001 |
| 0.5 | AVG | SVES+1 | SVES-3 | 0.6803 | 0.2230 | 0.2892 | 0.1937 | 0.0650 | 0.0129 | 0.2230 | 0.4255 | 0.0373 | 0.0129 |
| 0.5 | AVG | VES+1 | VES-3 | 0.1174 | 0.1174 | 0.2511 | 0.2511 | 0.0000 | 0.0044 | 0.1174 | 0.0049 | 0.0000 | 0.0125 |
| 0.5 | AVG-SF | ES+1 | ES-3 | 0.9007 | 0.2185 | 0.0000 | 0.0000 | 0.0008 | 0.0016 | 0.9007 | 0.0204 | 0.0000 | 0.0001 |
| 0.5 | AVG-SF | SVES+1 | SVES-3 | 0.2434 | 0.5375 | 0.0156 | 0.0002 | 0.0156 | 0.1424 | 0.1424 | 0.6336 | 0.0224 | 0.0129 |
| 0.5 | AVG-SF | VES+1 | VES-3 | 0.3094 | 0.0088 | 0.0000 | 0.0000 | 0.0300 | 0.0071 | 0.1608 | 0.0019 | 0.0000 | 0.0089 |
| 0.05 | AVG | ES+1 | ES-3 | 0.1686 | 0.3495 | 0.7807 | 0.9158 | 0.0041 | 0.0015 | 0.7807 | 0.0062 | 0.0000 | 0.0018 |
| 0.05 | AVG | SVES+1 | SVES-3 | 0.8777 | 0.2153 | 0.7567 | 0.6092 | 0.3483 | 0.0274 | 0.1790 | 0.8555 | 0.0873 | 0.0778 |
| 0.05 | AVG | VES+1 | VES-3 | 0.0718 | 0.9077 | 0.9077 | 0.4166 | 0.0044 | 0.0718 | 0.4166 | 0.0028 | 0.0001 | 0.0329 |
| 0.05 | AVG-SF | ES+1 | ES-3 | 0.8560 | 0.8856 | 0.0000 | 0.0000 | 0.0003 | 0.0647 | 0.8391 | 0.0016 | 0.0000 | 0.0011 |
| 0.05 | AVG-SF | SVES+1 | SVES-3 | 0.5182 | 0.4406 | 0.0094 | 0.0062 | 0.0094 | 0.4406 | 0.1269 | 0.5275 | 0.0524 | 0.0389 |
| 0.05 | AVG-SF | VES+1 | VES-3 | 0.2914 | 0.2264 | 0.0000 | 0.0000 | 0.0220 | 0.0652 | 0.3747 | 0.0007 | 0.0000 | 0.0220 |
| 0.001 | AVG | ES+1 | ES-3 | 0.1746 | 0.2937 | 0.5195 | 0.7507 | 0.0301 | 0.0022 | 0.4959 | 0.0027 | 0.0022 | 0.0095 |
| 0.001 | AVG | SVES+1 | SVES-3 | 0.8481 | 0.4013 | 0.8481 | 0.6424 | 0.5091 | 0.0290 | 0.1824 | 0.5091 | 0.1824 | 0.1613 |
| 0.001 | AVG | VES+1 | VES-3 | 0.0959 | 0.6559 | 0.6121 | 0.2316 | 0.0469 | 0.0959 | 0.6559 | 0.0020 | 0.0020 | 0.0959 |
| 0.001 | AVG-SF | ES+1 | ES-3 | 0.8809 | 0.9490 | 0.0000 | 0.0000 | 0.0006 | 0.5615 | 0.5615 | 0.0006 | 0.0006 | 0.0063 |
| 0.001 | AVG-SF | SVES+1 | SVES-3 | 0.6195 | 0.5869 | 0.0455 | 0.1252 | 0.0455 | 0.8537 | 0.1569 | 0.2976 | 0.1252 | 0.1075 |
| 0.001 | AVG-SF | VES+1 | VES-3 | 0.4158 | 0.5257 | 0.0000 | 0.0000 | 0.0306 | 0.3526 | 0.6103 | 0.0007 | 0.0009 | 0.0859 |

**Table 4:** *Multiverse Source Analysis of sources in the ES+1 condition in the* 130 ms *to* 200 ms *time window.*

| regu. parameter | agg. method | target condition | reference condition | Left Cingulate Gyrus positiveterior division | Right Cingulate Gyrus positiveterior division | Left Postcentral Gyrus | Right Postcentral Gyrus | Left Frontal Orbital Cortex | Right Frontal Orbital Cortex | Left Insular Cortex | Right Insular Cortex | Left Cingulate Gyrus anterior division | Right Cingulate Gyrus anterior division |
|---|---|---|---|---|---|---|---|---|---|---|---|---|---|
| 0.5 | AVG | VES | VES-3 | 0.1556 | 0.1945 | 0.1364 | 0.1556 | 0.0272 | 0.0120 | 0.0120 | 0.2310 | 0.5397 | 0.0120 |
| 0.5 | AVG-SF | VES | VES-3 | 0.1241 | 0.1805 | 0.0043 | 0.0082 | 0.0043 | 0.0073 | 0.0043 | 0.1805 | 0.5397 | 0.0043 |
| 0.05 | AVG | VES | VES-3 | 0.8894 | 0.8894 | 0.0076 | 0.0570 | 0.1667 | 0.0570 | 0.0076 | 0.0958 | 0.8894 | 0.0549 |
| 0.05 | AVG-SF | VES | VES-3 | 0.9027 | 0.9027 | 0.0846 | 0.1883 | 0.0846 | 0.0846 | 0.0047 | 0.0846 | 0.9027 | 0.0823 |
| 0.001 | AVG | VES | VES-3 | 0.9614 | 0.9614 | 0.0033 | 0.1360 | 0.3422 | 0.2249 | 0.0033 | 0.1360 | 0.9614 | 0.2441 |
| 0.001 | AVG-SF | VES | VES-3 | 0.8323 | 0.8310 | 0.2441 | 0.6735 | 0.2441 | 0.2441 | 0.0053 | 0.2296 | 0.9614 | 0.2441 |

**Table 5:** *Multiverse Source Analysis of sources in the VES condition in the* 220 ms *to* 330 ms *time window.*

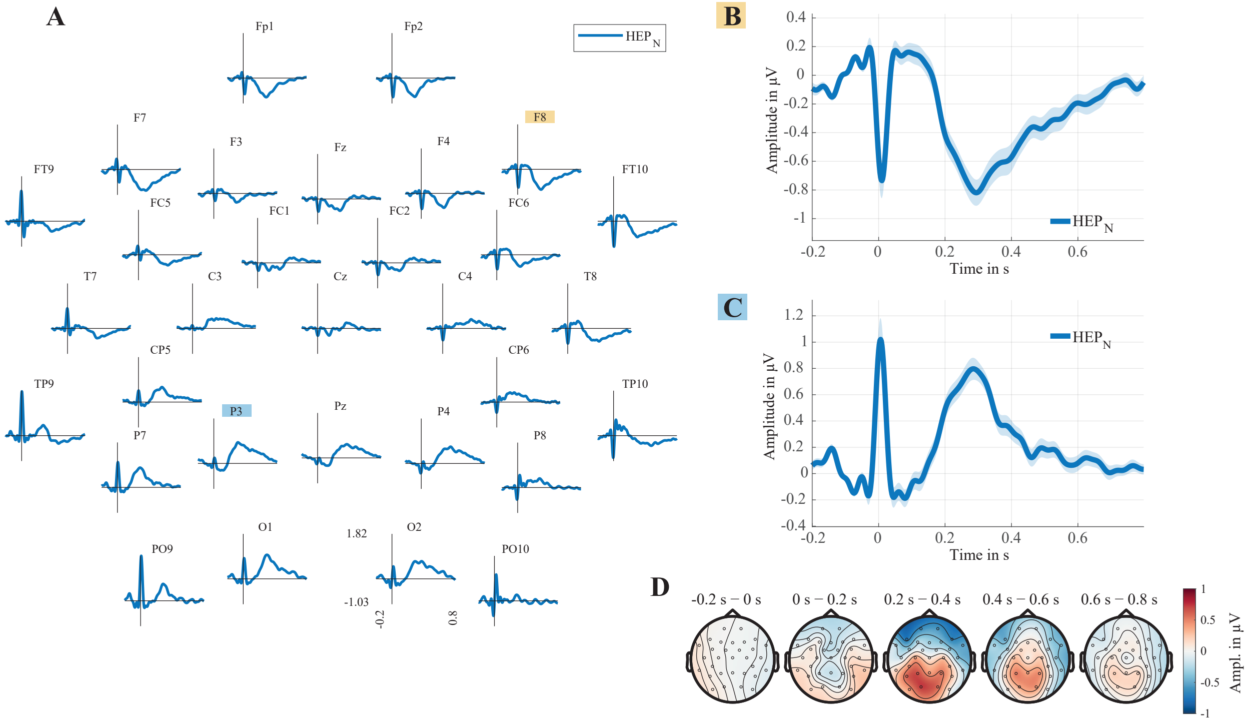

**Figure 15:** *HEP of sinus beats (N). Grand average over all participants.* **(A)** *HEP in each EEG channel.* **(B)** *HEP in electrode F8. The coloured-shaded regions around the lines mark the* ±1 *SEM.* **(C)** *HEP in electrode P3. The coloured-shaded regions around the lines mark the* ±1 *SEM.* **(D)** *Topographies over time.*

**Table 6:** *Matching of T-peak amplitude. Column Trials describes the number of remaining epochs per participant after matching.*

| | | Unmatched T-peak ampl. in µV | | Matched T-peak ampl. in µV | |
|---|---|---|---|---|---|
| | **Trials** | **ES-3** | **ES+1** | **ES-3** | **ES+1** |
| **Mean** | 29.10 | 166.81 | 159.38 | 159.79 | 159.18 |
| **Std** | 14.71 | 74.06 | 69.10 | 69.30 | 69.17 |
| **Min** | 8.00 | 38.73 | 28.95 | 29.08 | 28.95 |
| **Max** | 88.00 | 378.97 | 369.10 | 369.13 | 369.10 |
| $p$ | | 0.00 | | 0.77 | |

**Table 7:** *Matching of IBI. Column Trials describes the number of remaining epochs per participant after matching.*

| | | Unmatched IBI in ms | | Matched IBI in ms | |
|---|---|---|---|---|---|
| | **Trials** | **ES-3** | **ES+1** | **ES-3** | **ES+1** |
| **Mean** | 29.04 | 927.55 | 935.83 | 931.79 | 932.07 |
| **Std** | 14.85 | 133.07 | 138.78 | 135.05 | 135.27 |
| **Min** | 8.00 | 587.78 | 593.09 | 593.09 | 593.09 |
| **Max** | 105.00 | 1240.63 | 1264.21 | 1259.79 | 1261.05 |
| $p$ | | 0.00 | | 0.95 | |

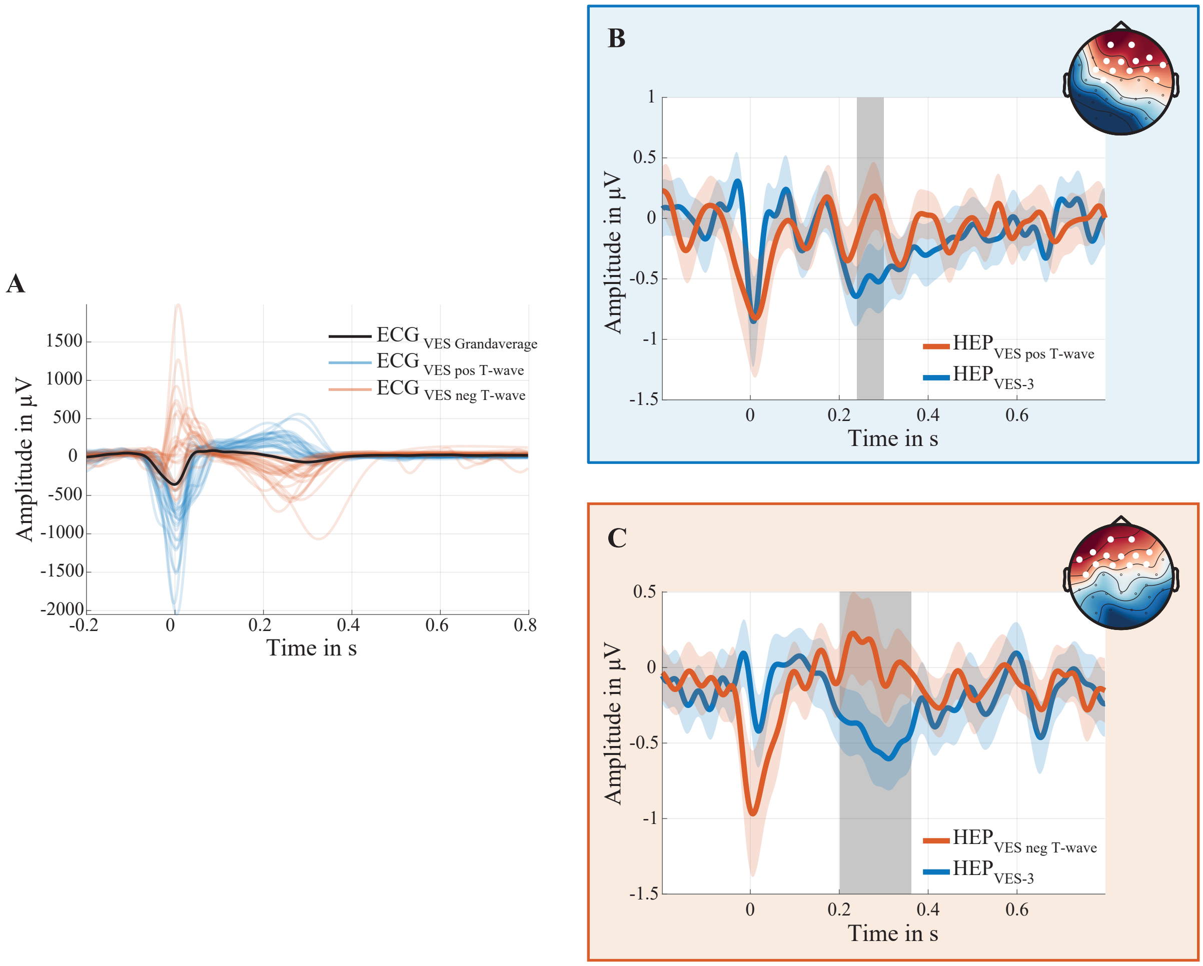


**Figure 16:** *Control Analysis for CFA of polymorph VES.* **(A)** *Polymorph appearance of VES. Blue lines represent participantwise averages of VES with positive T-wave and orange lines with negative T-waves respectively. Black line shows the grand average over all participants.* **(B)** *Significant difference between HEPs of subgroup VES with posivitve T-wave and reference condition ($p < 0.05$) averaged over the cluster of white marked electrodes.* **(C)** *Significant difference between HEPs of subgroup VES with negative T-wave and reference condition ($p < 0.05$) averaged over the cluster of white marked electrodes. The grey-shaded area highlights the time window of the found cluster. The coloured-shaded regions around the lines mark the ±1 SEM.*

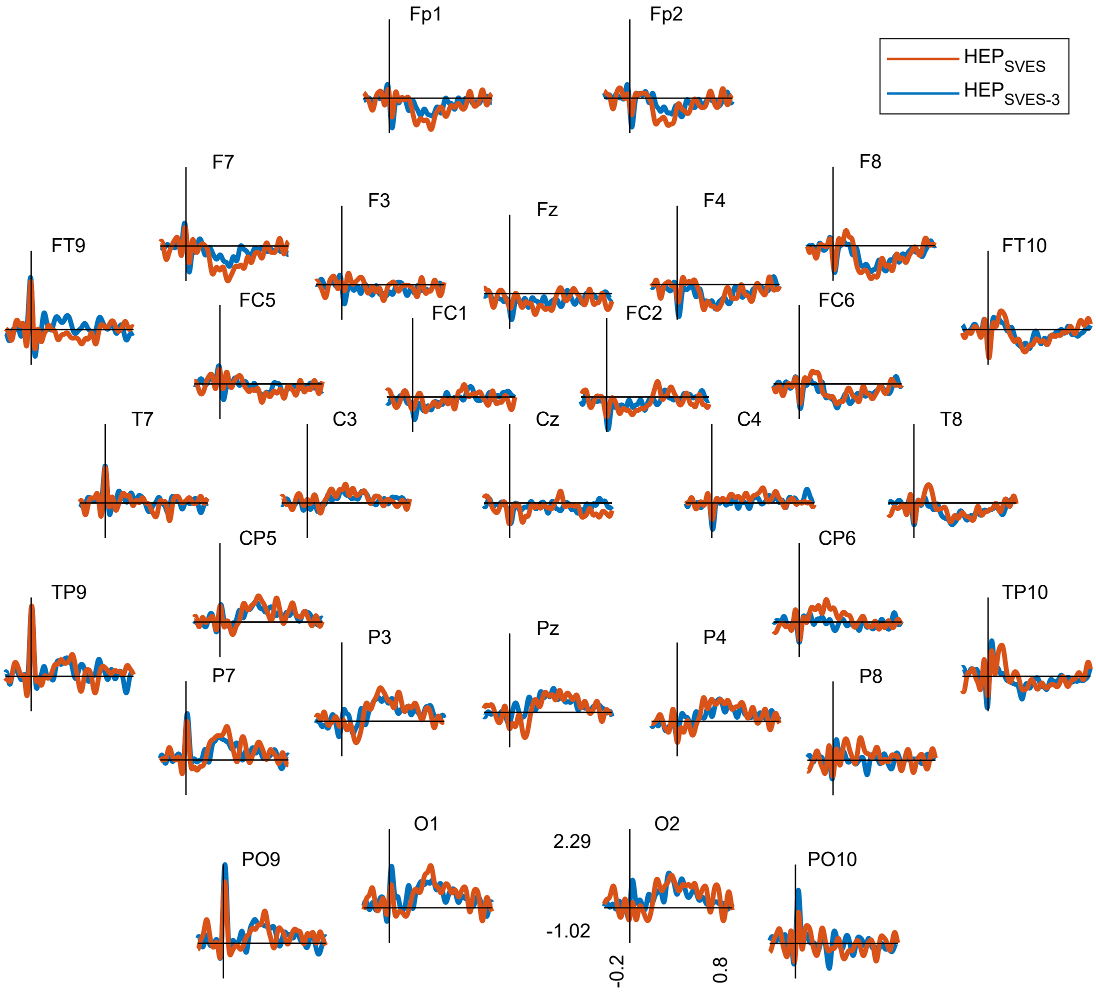


**Figure 17:** *No significant differences* ($p > 0.2$) *in HEP in the time domain between condition SVES and SVES-3.*

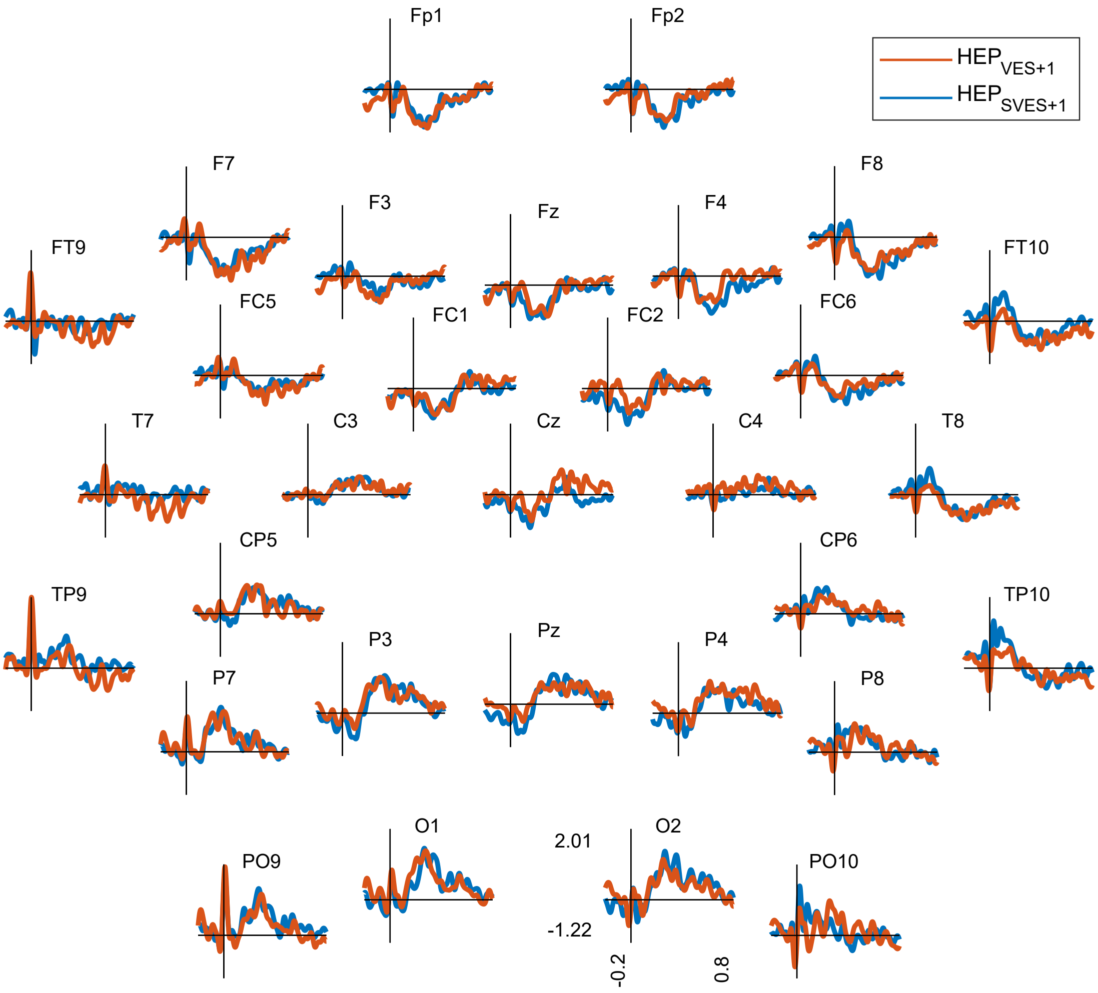


**Figure 18:** *No significant differences ($p > 0.5$) in HEP in the time domain between groups VES+1 and SVES+1.*

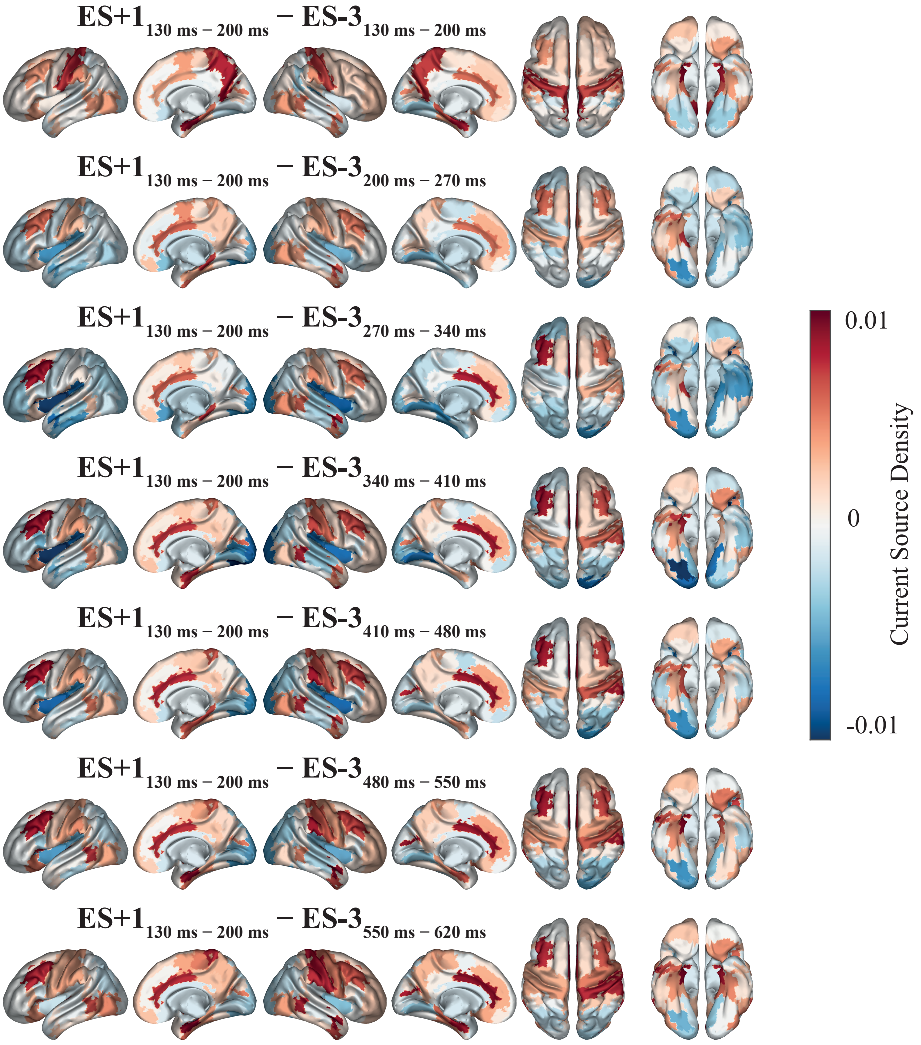


**Figure 19:** *Source Space of HEP difference between constant timewindow of ES+1 condition and moving time window across HEP of ES-3 condition. The ACC is across all comparisons significantly more active in the ES+1 condition ($p_{FDR} < 0.05$).*

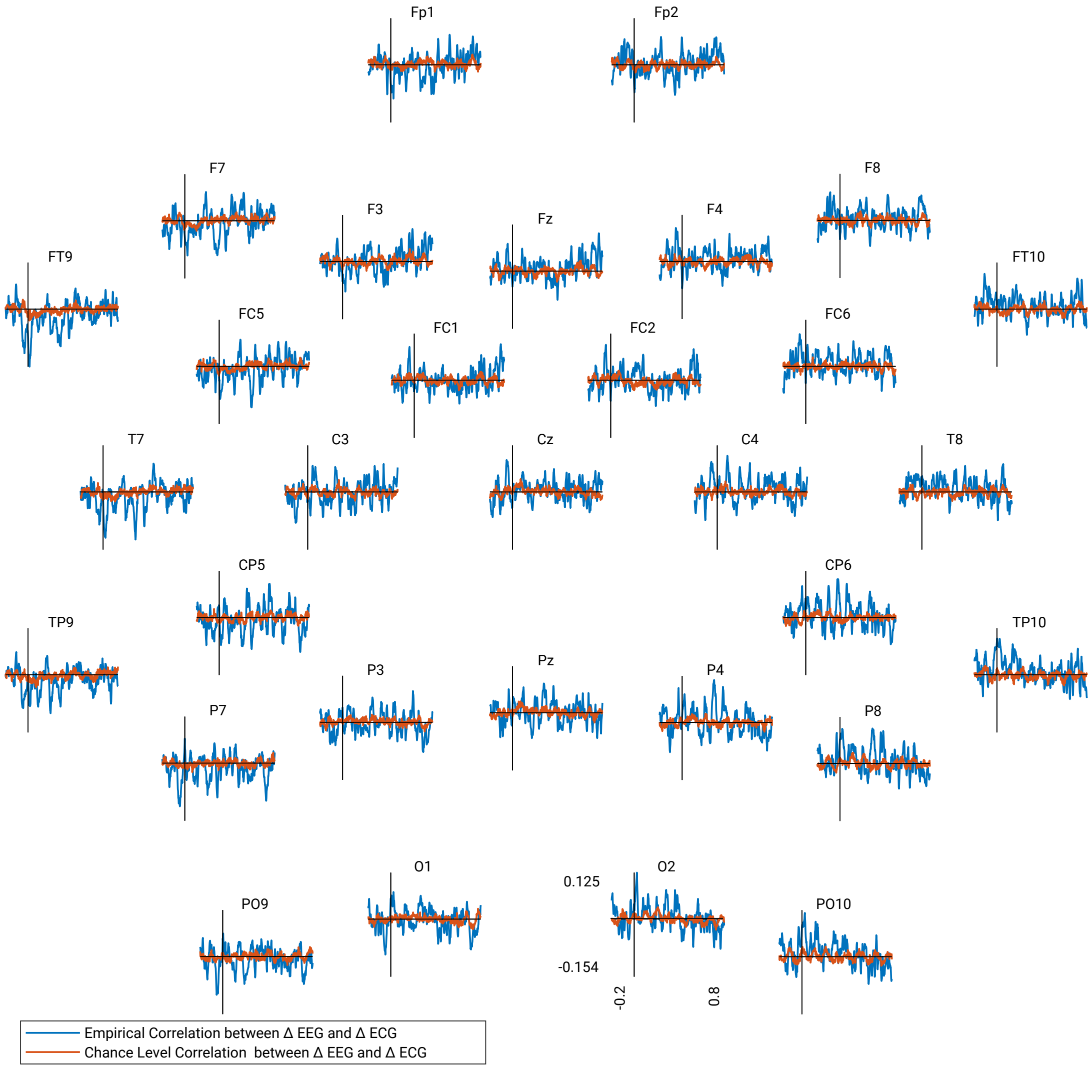


**Figure 20:** *Time course of the empirical and chance level fisher-transformed correlation between differences in the ECG and EEG for the VES with no significant findings.*

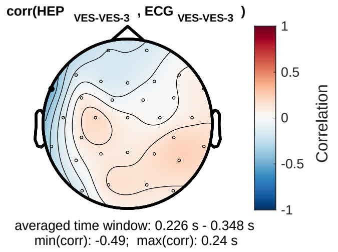


**Figure 21:** *Channelwise-correlation between differences in the ECG and EEG for the VES with a significant finding in electrode FT9 ($\rho = -0.49$; $p_{FDR} < 0.05$).*

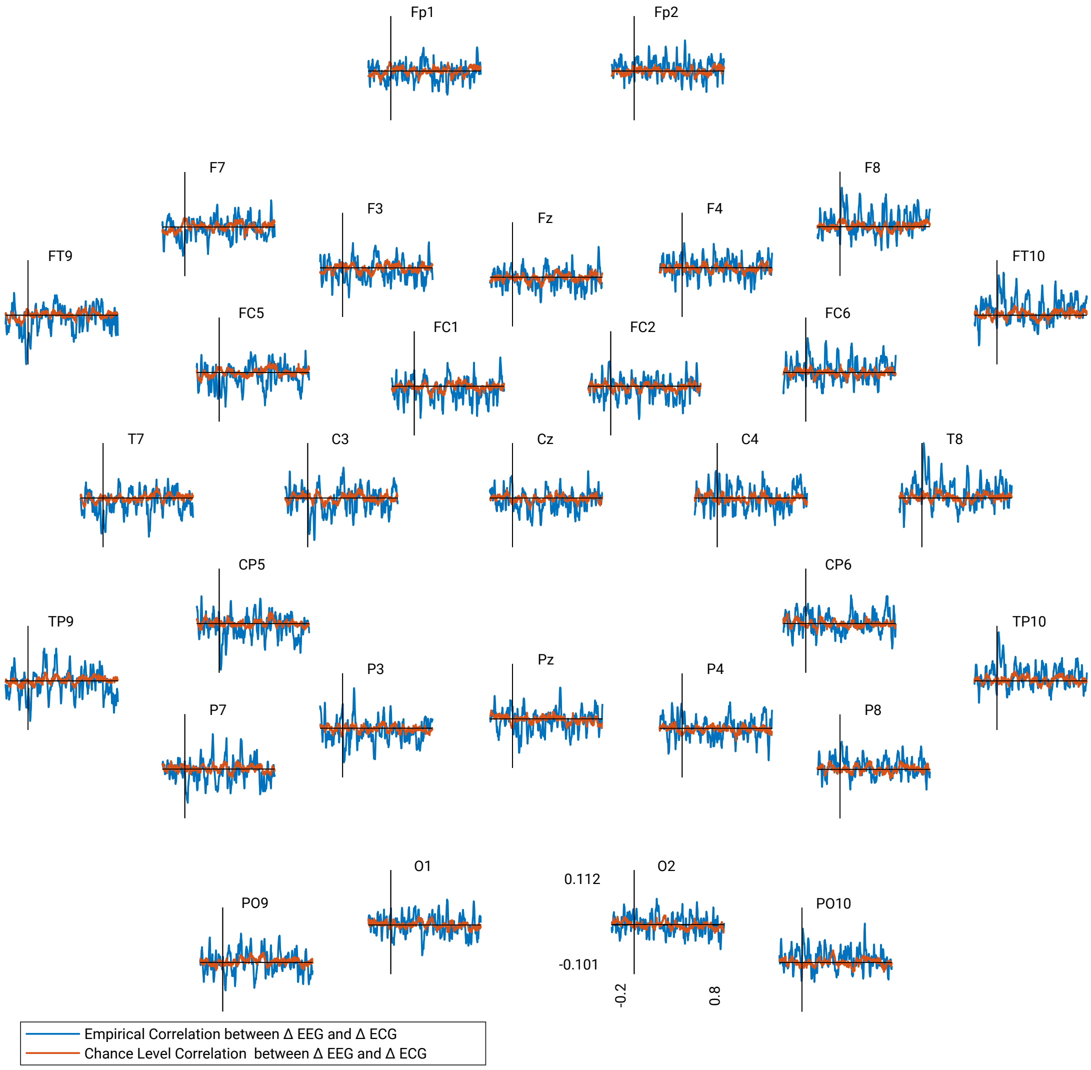


**Figure 22:** *Time course of the empirical and chance level fisher-transformed correlation between differences in the ECG and EEG for the ES+1 with no significant findings.*

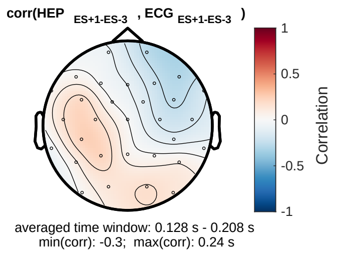


**Figure 23:** *Channelwise-correlation between differences in the ECG and EEG for the VES with no significant finding ($p_{FDR} > 0.05$).*

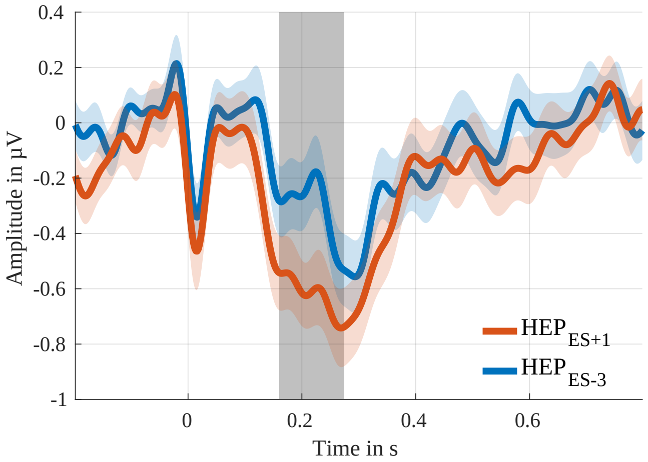


**Figure 24:** *Significant negative Cluster over central-frontal electrodes found in the time window between* 130 ms *to* 200 ms *(Monte Carlo* $p > 0.0942$*) between ES+1-condition and reference condition. The coloured-shaded areas around the lines mark the* ±1 *SEM.*

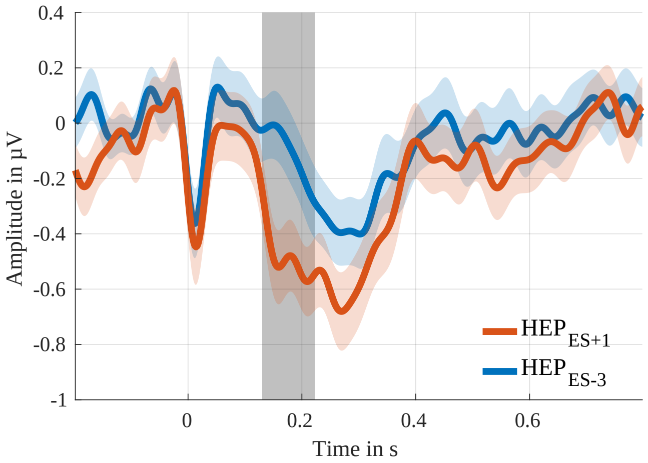


**Figure 25:** *Negative Cluster over central-frontal electrodes (Monte Carlo* $p = 0.002$*) between ES+1-condition and reference condition. The grey-shaded area highlights the time window of the found cluster. The coloured-shaded areas around the lines mark the* ±1 *SEM.*
